# Functional Role of Small Extrachromosomal Circular DNA in Colorectal Cancer

**DOI:** 10.1101/2025.08.01.668183

**Authors:** Judith Mary Hariprakash, Egija Zole, Weijia Feng, Dan Hao, Lasse Bøllehuus Hansen, Nirmalya Bandyopadhyay, Marghoob Mohiyuddin, Sihan Wu, Astrid Zedlitz Johansen, Julia Sidenius Johansen, Birgitte Regenberg

## Abstract

Extrachromosomal circular DNA (eccDNA) are circular DNA molecules that originate from chromosomal DNA but exist independently. While large eccDNA (ecDNA) contributes to tumorigenesis, the role of smaller eccDNA (<100,000 base pairs) in cancer remains unclear. Our analysis of 25 colorectal cancer (CRC) tumors and adjacent non-tumorous tissues revealed that eccDNA is significantly more abundant in tumor tissues, correlating strongly with chromosomal amplifications. The presence of whole intact genes on 1.29% of eccDNA was non-random. We identified 84 genes that recurred across tumors of multiple patients when present on eccDNA with 19% of genes being cancer-associated. eccDNA-borne genes were often accompanied by increased expression, and their contribution to expression was much larger than that from linear amplifications and the larger ecDNA. The cytokine gene CXCL5 exemplified this phenomenon, showing substantial copy-number increase and upregulation when present on eccDNA. Functional validation in cell lines showed that CXCL5 eccDNA enhanced transcriptional output and immune cell recruitment function. The recurrence and overexpression of CRC-related genes on eccDNA indicate their selection in tumors, suggest eccDNA can serve as a novel mechanism for dynamically influencing gene expression and is capable of conferring cancer phenotypes to cells. Analysis of chromatin landscapes revealed that eccDNA preferentially forms at sites of open chromatin and active transcription, with architectural boundaries marked by CTCF protein. Clinically, patients with higher eccDNA levels showed poorer relapse-free survival. These findings suggest that circular DNA elements across the entire size spectrum participate in cancer evolution, positioning eccDNA as a potential therapeutic target and prognostic biomarker.

## Introduction

Cancer is a complex disease defined by its remarkable variability across multiple dimensions including histological, clinical, cellular, molecular, and epigenetic (reviewed in (1, 2)). Among the emerging mechanisms contributing to tumor heterogeneity and adaptation, is extrachromosomal circular DNA (eccDNA) of chromosomal origin. These circular DNA molecules exist outside chromosomal DNA and range dramatically in size from less than a hundred base pairs (bp) to several megabases (3–5). The eccDNA family includes both small circles (<100 kb) from across the genome and larger, complex structures known as ecDNA (>100 kb) that often carry amplified oncogenes (3, 5–7). The role of ecDNA in driving tumor evolution through rapid oncogene amplification is well established (5, 8–11). This amplification can drive tumor heterogeneity and evolution by providing a mechanism for rapid gene copy number increase outside of chromosomal constraints. Consequently, ecDNA presumably contributes to the dynamic adaptability of cancer cells, influencing their growth, metastatic potential, and response to treatment (3, 5, 6, 9, 10, 12–14).

However, the population of smaller and more diverse circles eccDNA under 100,000 bp is poorly understood due to technical constraints (15). These circles originate from across the genome, especially from gene-rich chromosomes and repetitive sequence regions, and are found in both healthy somatic tissues and tumor tissues (3, 4, 11). Despite limited research, evidence suggests eccDNA may amplify oncogene copy numbers, generate novel oncogene isoforms, and serve as regulatory platforms for genetic adaptation (13, 15). The clinical relevance of eccDNA may be particularly apparent in colorectal cancer (CRC), one of the most prevalent malignancies in developed countries. Despite early detection through screening programs, CRC maintains high mortality rates and represents a significant health burden (16). Like many cancers, CRC exhibits substantial molecular and genetic heterogeneity (17–19). The disease progresses through dysregulation of key pathways: Wnt/β-catenin signaling drives early adenoma formation through APC mutations, RAS pathway mutations promote oncogenic survival signaling (reviewed in (20)) (21, 22), and PI3K/Akt/mTOR activation contributes to growth and treatment resistance (23). Additionally, chronic inflammation and immune evasion mechanisms facilitate CRC progression (24–26). For instance, CRC tumors often attract and activate immune cells, particularly neutrophils, through chemokines such as CXCL5 (27–29). Given this molecular complexity and the heterogenous nature of CRC, investigating the contribution of eccDNA in disease progression becomes particularly compelling. The current understanding of eccDNA in CRC is mostly limited to primarily cell lines (30, 31), larger ecDNA (32), and small patient cohorts (33). Nevertheless, a recent study has shown connection between eccDNA and colorectal cancer progression (34).

We hypothesize that eccDNA plays a critical role in CRC by facilitating increased expression of CRC-relevant genes with functional consequences for the tumors. We propose that such traits can lead to selection for CRC-relevant genes on eccDNA leading to their overrepresentation on eccDNA. Finally, we suggest that chromosomal focal amplifications contribute to increased levels of eccDNA within tumors with consequences for the patient. Using whole genome sequencing and Circle-Seq of matched tumor and adjacent normal tissues from CRC patients, we demonstrate that tumors harbor significantly more eccDNA than normal tissue. Patients with the highest eccDNA levels experience poorer relapse-free survival, suggesting prognostic value for eccDNA abundance. Importantly, eccDNA carrying complete oncogenes correlates with increased gene transcription in tumor tissue, confirming functional impact. The recurrent observation of specific genes on eccDNA across multiple patients indicates consistent oncogene selection patterns. These findings establish eccDNA not merely as a reflection of genomic instability, but as an active driver of oncogenic processes in colorectal cancer progression and evolution. Similar patterns of eccDNA-mediated oncogene amplification and poor clinical outcomes have been documented across diverse cancer types, including neuroblastoma with MYCN-containing eccDNA, hepatocellular carcinoma with miRNA-17-92 amplicons, and medulloblastoma, suggesting that eccDNA represents a fundamental mechanism of cancer evolution that transcends tissue-specific boundaries and warrant systematic investigation in colorectal cancer (3, 35, 36).

## Results

### eccDNA abundance is elevated in colorectal tumor tissue compared to non-tumorous adjacent tissue

To understand the landscape of circular DNA elements in colorectal cancer and investigate their effects on tumor phenotype, we performed circle-sequencing of eccDNA from tumor tissue (TT) and normal adjacent tissue (NAT) of 25 CRC patients removing both linear and mitochondrial DNA (37). RNA sequencing was performed on the matched TT and NAT samples to obtain corresponding transcriptome data. A subset of 12 TT samples was subjected to whole genome sequencing (WGS) to identify copy number variations (CNVs, **Figure 1a**). **Supplementary Table S1** provides the clinical characteristics of the sample cohort.

**Figure 1:**
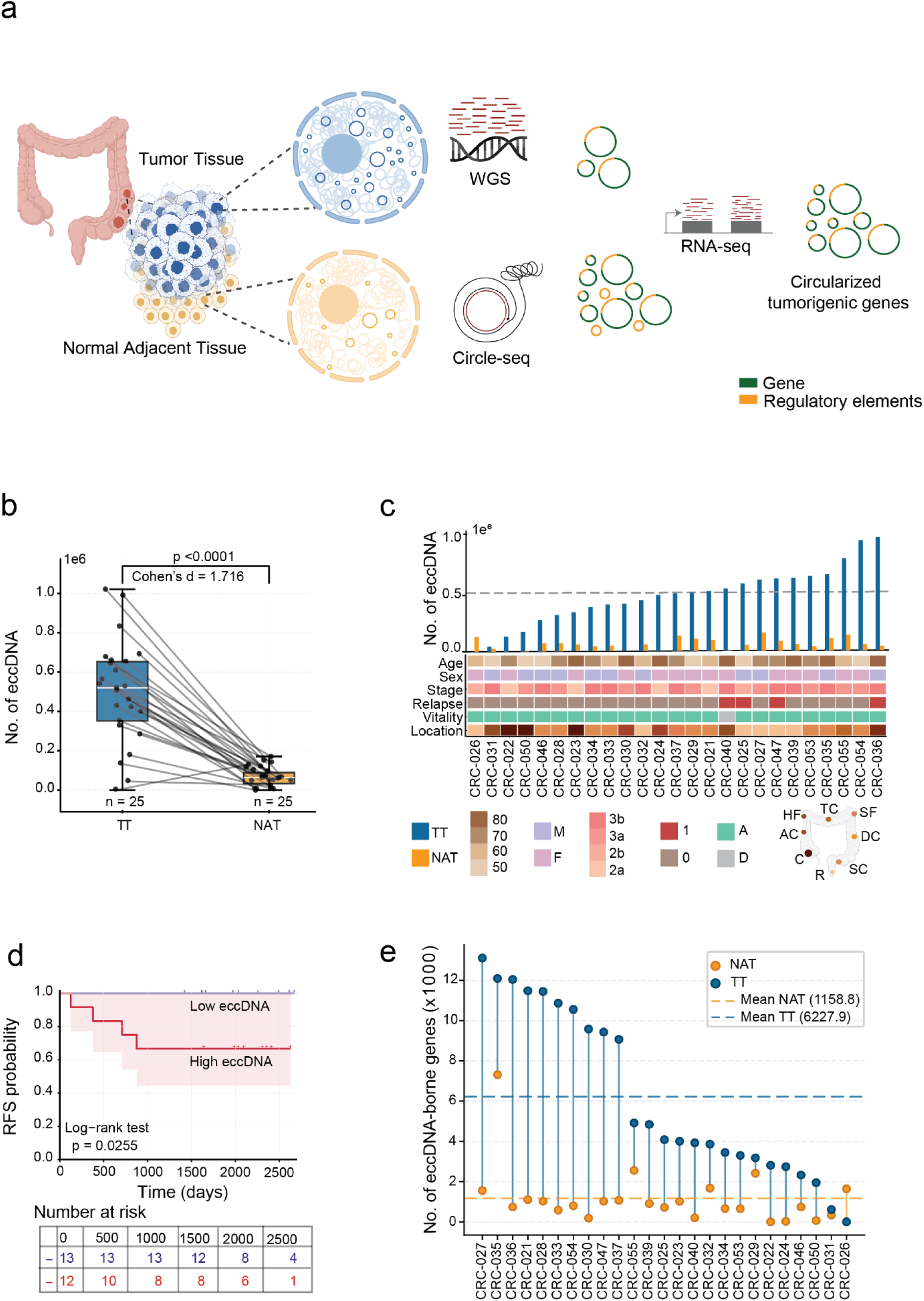
eccDNA in colorectal cancer. **(a)** Workflow for identification of circularized tumorigenic genes. Tumor tissue and matched normal adjacent tissue samples are collected from colorectal cancer patients. Genomic DNA is extracted and subjected to WGS to identify genetic alterations. Circle-seq methodology is employed to detect eccDNA elements. RNA-seq analysis is performed to characterize gene expression profiles. Integration of these multi-omics approaches enables the identification of circularized tumorigenic genes. (**b**)Quantitative comparison of eccDNA counts between NAT and TT samples. The boxplot displays the distribution of eccDNA counts in NAT (yellow, n = 25) and TT (blue, n = 25) samples. Each box represents the interquartile range (IQR), which contains the middle 50% of the data, with the horizontal line inside the box indicating the median. The whiskers extend to the minimum and maximum values within 1.5 times the IQR. Individual data points are plotted as dots, representing specific samples. Gray lines connect paired NAT and TT samples from the same patient. The exact p-value =2.58×10^-9^ derived from the Wilcoxon rank-sum test and Cohen’s d = 1.716. **(c)** Patient-specific eccDNA profiles and clinical characteristics. The bar plot shows the number of eccDNA molecules (y-axis, up to 1.0×10^6^) detected in TT (blue) and NAT (Yellow) samples for each patient (x-axis, ordered according to the total number of eccDNA in TT). Below the graph, a clinical annotation plot displays various characteristics for each patient, including age, sex, cancer stage, relapse status, vitality, and tumor location. **(d)** Kaplan-Meier survival curves comparing relapse-free survival probability between colorectal cancer patients with high (red line) and low (purple line) eccDNA levels, stratified based on the median eccDNA counts. The x-axis shows time in days (0 to 2800), and the y-axis represents the relapse-free survival probability (0 to 1.0). The difference in the survival curve is significant, with a p-value of 0.0255, as determined by the log-rank test. **(e)** Paired comparison of eccDNA-harboring complete genes between TT (blue) and NAT (orange) in colorectal cancer patients (n=25). Each connected pair represents one patient. Connecting lines indicate direction of change: blue lines show TT > NAT, orange lines show TT < NAT. Dashed lines represent mean values. Patients are ordered by decreasing TT eccDNA count.

Circle-seq analysis identified 14,242,467 eccDNA elements across 25 samples, with 12,460,376 in TT and 1,782,091 in NAT samples. Notably, one tumor sample and one normal adjacent tissue sample had no detectable eccDNA. We observed significantly higher numbers of eccDNA in TT compared to NAT samples (mean eccDNA count: TT = 519,182, NAT = 74,253; Wilcoxon rank-sum test, p = 2.45×10^-7^; **Figure 1b**). This represents a 7-fold increase in eccDNA abundance in tumor tissue, with a large effect size (Cohen’s d = 1.716). **Figure 1c** illustrates the eccDNA counts in paired tissues for each patient and their clinical characteristics. The eccDNA numbers did not correlate with the clinical characteristics of patients, such as age, sex, cancer stage, and tumor location (linear model, p-value = 0.7132). However, when we stratified patients into low (n=13) and high (n=12) eccDNA groups based on median tumor eccDNA count, patients in the low eccDNA group had no disease relapse (0/13, 0%) during the follow-up period, while 4 of 12 patients (33.3%) in the high eccDNA group experienced relapses. **Figure 1d** illustrates the Kaplan-Meier curves of the two groups showing significant difference in relapse-free survival with log-rank p=0.025. Due to the absence of events in the low eccDNA group, hazard ratio estimation was not feasible. The model showed good discriminative ability (C-index=0.789, 95% CI: 0.687-0.891).

### eccDNA harbors intact genes

We next aimed to determine if eccDNA serves a role beyond a structural genomic element by expressing intact genes within eccDNA (hereafter referred to as ’eccDNA-borne genes’). We observed that the number of eccDNA-borne genes per TT sample ranged from 647 to 16,184, with an average of 7,884 (**Figure 1e**). Notably, 1.29% on average harbored complete intact genes in tumor samples.

To assess whether complete genes were overrepresented on eccDNA, we compared the observed number of intact genes within eccDNA to expectations from 25 random genomic region datasets matched for size distribution and count per sample. This analysis revealed a burden-dependent pattern: samples with lower total gene counts (log2(counts+1) < 3.5) showed enrichment of complete genes within eccDNA compared to random expectation, while samples with higher total gene counts (log2(counts+1) > 3.5) showed depletion, with random regions containing more genes than eccDNA (**Supplementary Figure 1a**). Statistical analysis using permutation testing (n = 10,000) revealed that 20 of 24 samples showed significant deviation from random expectation (p < 0.05) (**Supplementary Table S2**).

To ascertain whether the increased eccDNA-borne genes could be attributed to higher gene density, we analyzed any potential correlation between gene density and eccDNA-borne genes per chromosome. We observed no correlation (linear regression, R^2^= -0.240, -0.095 for TT and NAT, respectively) between gene density and eccDNA-borne genes (**Supplementary Figure 1b**). These results suggest that eccDNA is a significant repository of genetic information and that factors other than gene density may contribute to the generation or retention of eccDNA within the genome.

### Genes on circular DNA exhibit higher expression than linear DNA counterparts

Previous studies have shown that genes on circular DNA >100,000 bp (ecDNA) are often highly overexpressed in many tumors (3, 5, 7). To compare the contributions from eccDNA, ecDNA, and linear amplification to transcription, we employed Amplicon Architect to analyze the WGS data of 12 TT samples, identifying ecDNA and linearly amplified genes. This analysis identified 346 linearly amplified genes and 19 ecDNA-borne genes across the samples, in addition to 34,850 ecDNA-borne genes identified by the Circle-Seq analysis in the same samples. We then compared the expression levels of genes in linear DNA, ecDNA, and eccDNA using z-scores. Notably, while ecDNA-borne genes displayed the highest median expression levels, eccDNA contributed most of highly expressed circular genes overall (n=38,107 eccDNA genes versus n=19 ecDNA genes), with many eccDNA-borne genes reaching high z-scores in the upper tail of the distribution (**Figure 2a**). Genes within eccDNA and ecDNA exhibited significantly higher expression levels than those in linear DNA (Kruskal-Wallis test p = 5×10^-07^). This finding underscores the circular DNA elements across the entire size spectrum enhances gene expression. We further examined the diversity and complexity of eccDNA-borne genes by analyzing their lengths and gene categories. Gene lengths ranged from 8 bp (TRDD1) to 2,473,620 bp (RBFOX1), with a mean size of 21,291 bp (**Supplementary Figure 1c**). For protein-coding genes specifically, the size distribution (median: 25,554 bp; mean: 67,087 bp) closely resembled that of protein-coding genes across the genome (median: 27,411 bp; mean: 68,851 bp), suggesting no substantial size bias in eccDNA gene capture. In terms of gene categories, pseudogenes, protein-coding genes, snRNAs, and lncRNAs were the most frequently observed among eccDNA-borne genes (**Figure 2b**).

**Figure 2:**
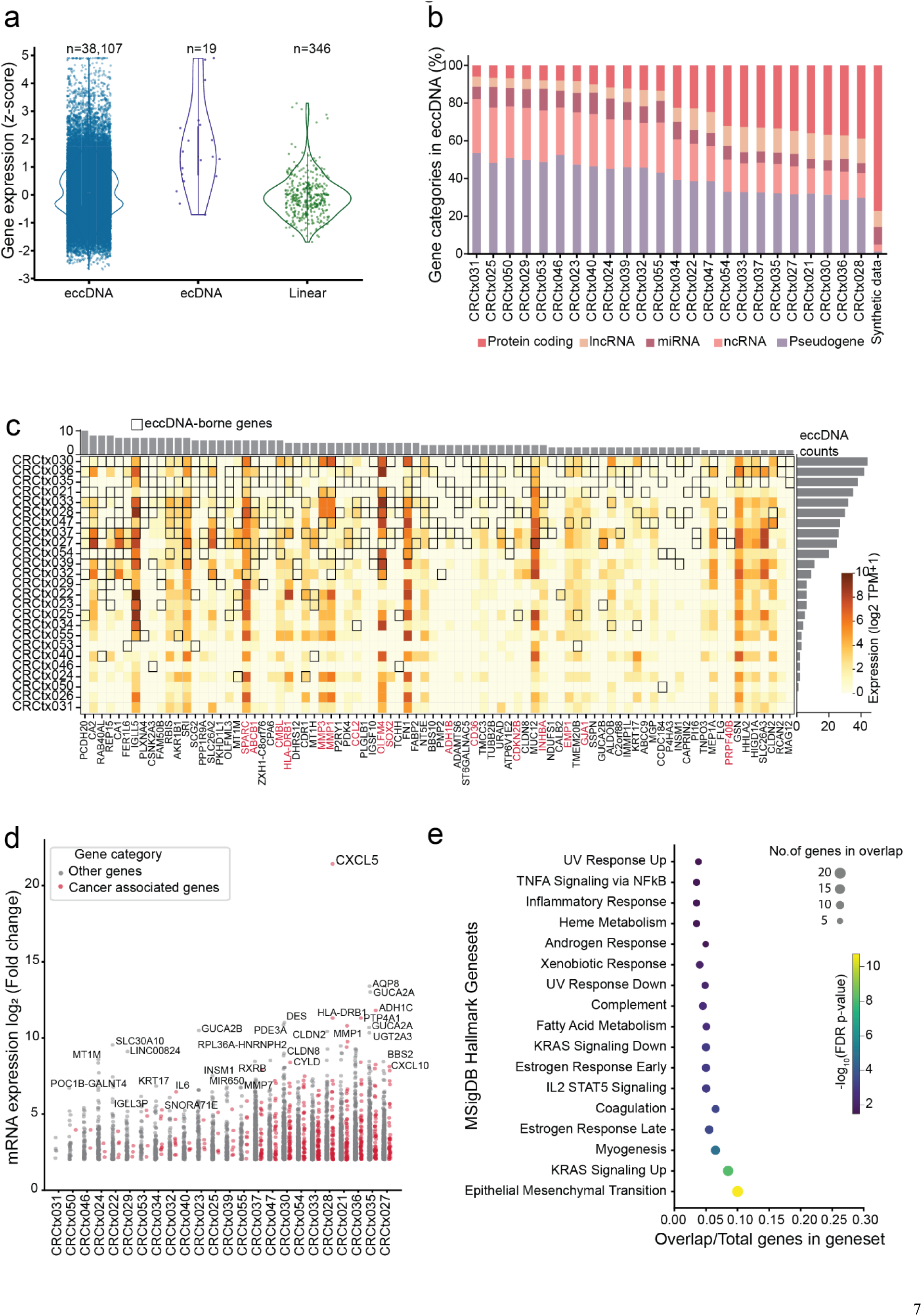
Characterization of eccDNA containing genes in colorectal cancer samples. **(a)** Violin plots comparing the expression levels (z-score) of genes associated with different forms: eccDNA (identified via circle-seq), ecDNA (identified via amplicon architect), and linear RNAs. The y-axis represents the z-scored gene expression levels, showing the distribution and density of expression values across these groups. The number of genes identified in each category is at the top of each violin plot. **(b)** A stacked bar chart shows the gene categories of eccDNA harboring complete genes in each sample. Different colors represent various gene categories based on RefSeq annotations, including protein-coding genes, lncRNAs, miRNAs, pseudogenes, and ncRNAs. **(c)** Heatmap displaying the frequency and expression of eccDNA-borne genes across samples. The color gradient indicates log_2_ gene expression levels in TPM (transcripts per million), while black boxes highlight the eccDNA-borne gene in the sample. The bar plots to the right and top show the total counts of eccDNA-borne genes. Gene names in red indicate cancer-associated genes. **(d)** Scatter plot showing the fold-change in expression of eccDNA-borne genes in TT relative to NAT. Red dots indicate cancer-associated genes, while grey dots represent other genes. The top 30 gene symbols are labeled in the figure. (**e**) The scatter plot shows the over-representation of differentially expressed eccDNA-borne genes in the MSigDB Hallmark gene set. The x-axis represents the ratio of the overlapping genes to the total number of genes in the pathway. The size of the circle denotes the number of genes in overlap, and the color shows the negative logarithmic adjusted p-value.

### eccDNA-borne genes are associated with carcinogenesis and show enhanced expression

Having established the high expression of eccDNA-borne genes, we investigated if certain genes on eccDNA had traits of selection by focusing on recurrent genes and their expression patterns in colorectal cancer. We first examined gene recurrence across our 25 tumor samples and identified 84 eccDNA-borne genes that appeared in ≥5 samples (∼20% of samples) (**Figure 2c**). Among which, 19% (16/84) were cancer-associated, including well-characterized oncogenes such as *SPARC, ABCB1, CMBL, HLA-DRB1, MMP3, MMP1, CCL2, OLFM4,* and *SOX2*.

To assess functional relevance, we then examined expression changes in eccDNA-borne genes between paired TT and NAT samples. We identified 1,480 eccDNA-borne genes that were differentially expressed (|log₂ fold change| > 1, p ≤ 0.05) between TT and NAT (**Figure 2d**). Notably, 15.1% (223/1480) were involved in known cancer-related functions, significantly more than the expected background proportion of 10.0% based on the total transcriptome (Binomial test, p = 6.4×10⁻¹⁰; **Supplementary Table S3**).

To further examine if eccDNA borne genes were selected in the tumor tissue, we performed gene-set enrichment analysis on the top 500 upregulated eccDNA-borne genes using two gene sets from the MSigDB database: (i) Hallmark and (ii) C6 oncogenic gene sets. Hallmark gene set analysis revealed a significant enrichment of cancer associated pathways (FDR corrected p <0.01), including epithelial-mesenchymal transition (EMT), KRAS signaling, immune responses, and metabolism (**Figure 2e**). The overlap between genes in each hallmark set and our upregulated gene list ranged from approximately 5% to 30%, with the EMT showing the highest overlap percentage. We found several significantly enriched c6 oncogenic gene sets as well, with a particular emphasis on twelve KRAS-related gene sets, along with gene sets associated with P53, PTEN, IL2, WNT signaling, and ATF2 pathways (**Supplementary Figure 1d, Supplementary Table S3**).

A striking example of eccDNA-driven transcriptional activation was observed in the *C-X-C motif chemokine ligand 5* (CXCL5) gene (**Figure 2d**). Tumor sample CRCtx028 contained an eccDNA (size <100 kb) harboring CXCL5 and gene expression showed a dramatic ∼2²¹-fold increase compared to paired NAT. This exemplifies how even small eccDNA elements can drive profound transcriptional upregulation of cancer-relevant genes. Our analysis suggests that eccDNA-borne genes significantly influence gene expression in colorectal cancer, with recurrent oncogenes and differential expression patterns indicating a selective advantage in tumor progression. The enrichment of cancer-related pathways, particularly KRAS signaling, and EMT, coupled with dramatic examples of gene upregulation like CXCL5, underscores the functional importance of eccDNA in tumor biology.

### Functional validation demonstrates eccDNA-mediated phenotypic effects

To investigate the potential direct association between eccDNA-borne genes and tumorigenesis, we focused on the CXCL5 gene identified on eccDNA in the CRCtx028 colorectal cancer sample (**Figure 3a**). We observed CXCL5 to be expressed only in the TT sample, while no expression was recorded in the NAT sample. To confirm the presence and structure of the CXCL5 eccDNA (hereafter referred as [CXCL5*^circle^*]), we confirmed the junction site by PCR on agarose gel in the CRCtx028 sample for one possible circle (**Figure 3b**). Quantitative PCR revealed that the copy number of the CXCL5 gene was up to 20 times more in the tumor sample, while no CXCL5 gene was found in the control NAT, after removing linear DNA (**Figure 3c**) (**Supplementary Figure 2a, b**). These results suggest amplification of CXCL5 via eccDNA in the tumor tissue. To further understand the origin and mechanism of this overexpression, we attempted to identify allele-specific variants that could link the eccDNA to its chromosomal source through haplotyping. However, we did not observe any allele-specific markers on the eccDNA that would allow us to determine the haplotype from which it originated. This limitation prevents us from definitively connecting the observed CXCL5 expression to a specific chromosomal allele.

**Figure 3:**
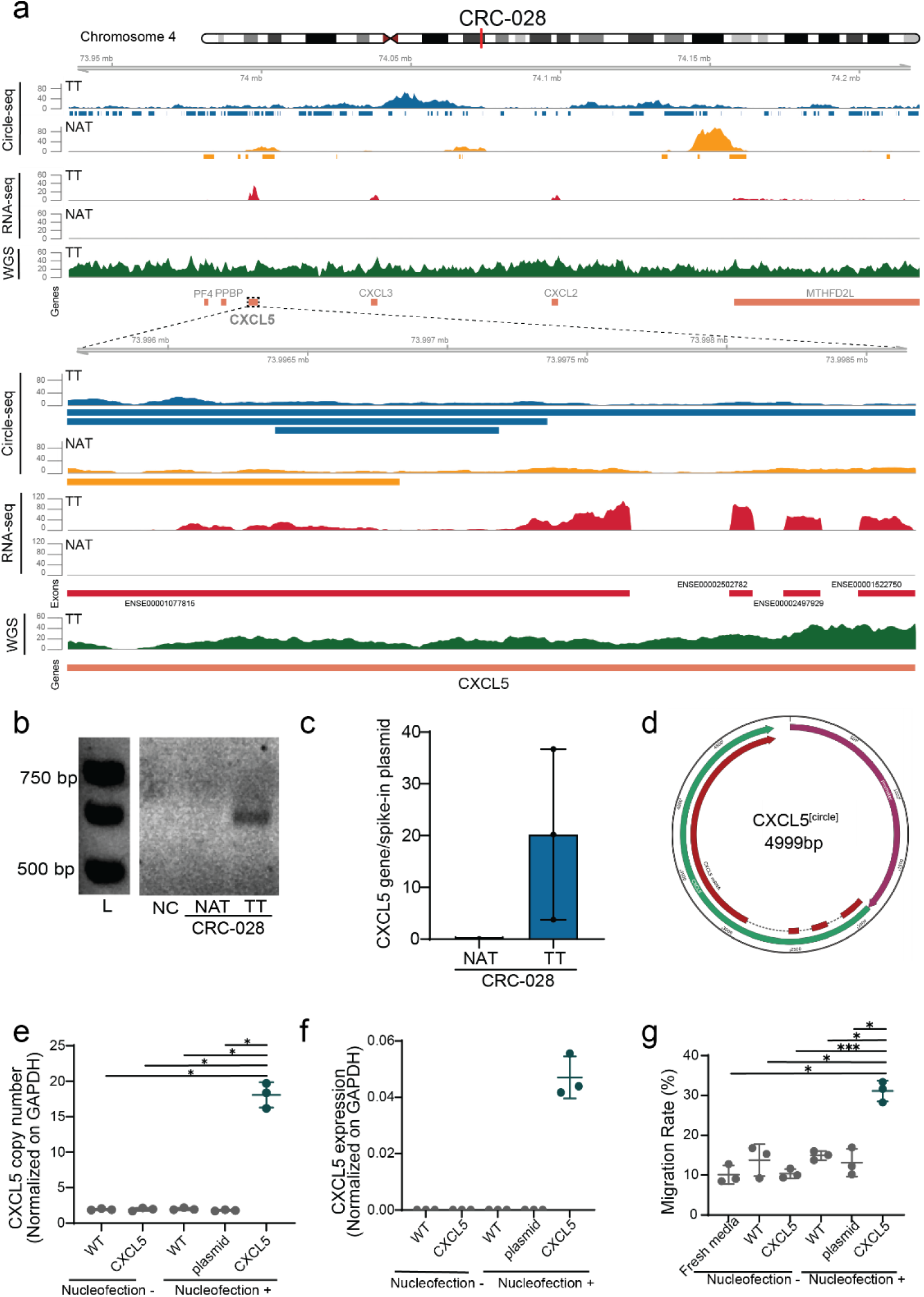
CXCL5 circle expression. **(a)** Genomic and transcriptomic profile of the CXCL5 locus in colorectal cancer patient CRC-028. The figure shows various data tracks aligned to a section of chromosome 4 (73.5Mb - 74.2Mb). Top: Ideogram of chromosome 4 with the analyzed region highlighted. eccDNA: Blue and yellow tracks show eccDNA coverage from circle-seq in the TT and NAT, respectively. RNA-seq: Red tracks depict RNA-seq read coverage, reflecting gene expression levels in TT and NAT samples. WGS: Green track represents whole genome sequencing (WGS) coverage, showing genomic read depth. Genes: Gene annotations for the region, with CXCL5 highlighted. Bottom: The same data tracks are displayed in a zoomed-in view of the CXCL5 locus (73.996 Mb—73.998Mb) eccDNA: Detailed eccDNA coverage at the CXCL5 locus for TT (blue) and NAT (yellow). RNA-seq: Higher resolution RNA-seq read coverage, highlighting differential gene expression of CXCL5 between TT and NAT samples. WGS: Zoomed-in WGS coverage at the CXCL5 locus, illustrating genomic read depth. Genes: Detailed annotations of the CXCL5 gene, including exons, are displayed. **(b)** PCR amplicon products of the native [CXCL5^circle^] junction sites in CRC-028 patient samples on an agarose gel. L-1 kb ladder, NC-negative control. **(c)** CNV of CXCL5 gene on eccDNA in CRC-028 patient samples measured by qPCR assay. The circular p4339 plasmid, an internal control, was used to normalize CXCL5 CN among the samples. **(d)**Map of the [CXCL5*^circle^*] construct. Circular map of the 4,999 base pair (bp) CXCL5 eccDNA construct. The circular structure contains the following key elements: the promoter region and the CXCL5 mRNA region. Gray arrows indicate the direction of the transcription, and dashed lines represent intron regions. Base pair positions are marked around the circumference in 500 bp intervals from 500 to 4500. **(e)** CXCL5 CNV following transfection with one µg of artificial circular [CXCL5^circle^] DNA in colorectal cell line SW620 compared to control conditions (f-one-way anova, p = 6×10^-10^; *, *** indicates Tukey HSD, family-wise error rate <0.05 and 0.001 respectively), assessed by droplet digital PCR (ddPCR). A plasmid containing a random sequence was used as a negative eccDNA control. WT-wild type. **(f)** CXCL5 RNA expression following the artificial [CXCL5*^circle^*] transfection in comparison to control conditions, measured by ddPCR. **(g)** Boyden Chamber cell migration assay for non-transfected cell line THP1, using spent medium from the transfected SW620 cells with the [CXCL5*^circle^*] (f-one-way anova, p = 5×10^-06^; * indicates Tukey HSD, family-wise error rate <0.05)

Given these limitations in identifying the chromosomal origin of the natural CXCL5 eccDNA, we decided to take a synthetic approach to directly test the functional implications of CXCL5 eccDNA in colorectal cancer (**Supplementary Figure 2c**). To achieve this, we developed a synthetic [*CXCL5^circle^*] (**Figure 3d**) construct containing the 3,035 bp long CXCL5 gene and its 1,845 bp promoter and transfected it into the colon cancer cell line SW620. Transfection of one million SW620 cells with 1.0 µg [CXCL5*^circle^*] led to an average copy number of 18.07±1.45 CXCL5, while SW620 cells transfected with an eccDNA containing a random sequence only carried the expected two copies of CXCL5 like the parental SW620 control (ANOVA F = 240.04, p = 6.92×10^-10^; Tukey’s HSD (honestly significant difference) for CXCL5_pos vs plasmid_pos p < 0.001,# and CXCL5_pos vs WT_neg p < 0.001) (**Figure 3e**). Cells exposed to the [CXCL5*^circle^*] without transfection did not increase the copy number of CXCL5, indicating that transfection successfully caused the [CXCL5*^circle^*] to enter SW620. We found that the transfected [CXCL5*^circle^*] was expressed in the SW620 cell line, while no expression of the CXCL5 gene was detected in any of the controls (**Figure 3f**).

The expression of CXCL5 was also reflected at the phenotypic level. The *CXCL5* chemokine is known to induce migration of monocytes and other immune cells (38). We set up a Boyden Chamber cell migration assay in which THP1 monocytes were exposed to medium from SW620 with and without [CXCL5*^circle^*] and tested for their ability to migrate through a polycarbonate membrane. We found that spent medium from SW620 [CXCL5*^circle^*] cells induced a significantly higher migration rate of the monocyte cell line THP1 than spent medium from SW620 cells not transfected with [CXCL5*^circle^*] (**Figure 3g**) (ANOVA F = 25.74, p = 5.04×10^-06^; Tukey’s HSD CXCL5_pos vs CXCL5_neg p < 0.001, mean difference = 20.75). This enhanced migration was specific to the CXCL5 eccDNA, as no significant differences were observed between control conditions including untransfected cells and cells transfected with control plasmid (Tukey’s HSD p > 0.05 for all control comparisons). In short, these experiments reveal that eccDNA in CRC is associated with increased expression of genes, and that expression of genes from synthetic eccDNA can allow cell lines to express cancer phenotypes. The ability of eccDNA to activate the CXCL5 gene expression and influence immune cell recruitment further corroborates that eccDNA of all sizes can serve as a mechanism for dynamically controlling gene expression and that eccDNA has the potential to confer cells cancer phenotypes.

### eccDNA exhibits size periodicity, chromosomal bias, and chromatin boundary associations

We examined eccDNA size distribution across our samples, revealing distinct patterns between tumor and normal tissues. The median size of circles was 1,279 bp and 1,842 bp in TT and NAT, respectively. Furthermore, around 85% of the eccDNA detected were less than 2 kilobase (kb), and signatures of eccDNA were found in sizes up to 32 megabases (Mb) (**Supplementary Figure 3a**).

The size distribution of eccDNA exhibited strong nucleosome periodicity, particularly in tumor samples (**Figure 4a**). The most prominent peak was observed at 360 bp corresponding to the length of 2-nucleosome (2N), containing 10.9% of tumor eccDNA compared to only 3.4% in NAT (3.2-fold enrichment, Chi-square test p < 0.0001). Secondary peaks at 3N (540 bp) and 4N (720 bp) positions showed 1.8-fold enrichments (both p < 0.0001). Overall, 40.2% of TT eccDNA aligned with nucleosome boundaries compared to 34.6% in NAT.

**Figure 4:**
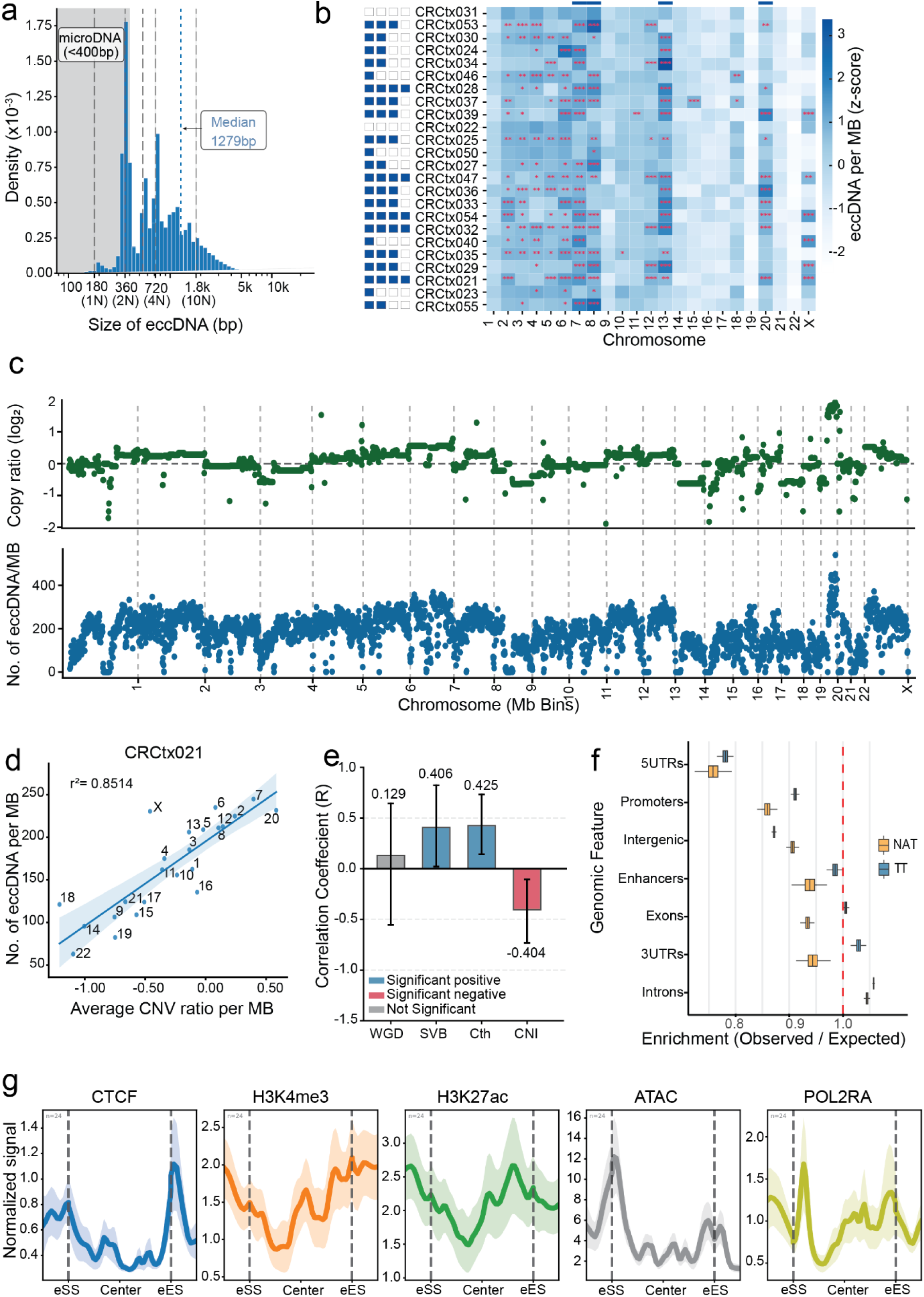
Genomic and epigenomic characteristics of eccDNA in colorectal cancer. **(a)** Density plot illustrates the size distribution of eccDNA in TT relative to nucleosome units. The gray area represents microDNA. The vertical blue dashed line indicates the median eccDNA size (1279 bp). The x-axis represents fragment sizes ranging from 1N to approximately 68N (10 kb) nucleosome units, where 1N corresponds to 147 bp of DNA wrapped around a single nucleosome core, displayed on a logarithmic scale. The y-axis denotes the density of eccDNA occurrences in the size range. **(b)** Heatmap illustrates the genomic distribution of eccDNA across chromosomes for individual patients. The y-axis lists patient identifiers (CRCtx031 to CRCtx055), while the x-axis represents chromosomes (chr1 to chr22, chrX). The color intensity indicates the relative abundance of eccDNA per Mb (z-score), with darker blue representing higher abundance and lighter shades indicating lower abundance. Red asterisks denote statistical significance levels: * p<0.05, ** p<0.01, *** p<0.001. Blue boxes highlight samples with significant enrichments in chromosomes 7, 8, 13, and 20, which represent recurrent hotspots of eccDNA formation across multiple patients. **(c)** Genome-wide distribution of eccDNA and copy number for sample CRCtx021. The top panel (green) shows the log_2_ copy number ratio across chromosomes per MB obtained from CNVKit, while the bottom panel (blue) represents the corresponding eccDNA counts per MB. **(d)** Correlation between eccDNA counts per MB and average copy number across chromosomes in colorectal cancer sample CRCtx021. Each point represents an individual chromosome. The blue line indicates the linear regression fit, with the shaded area showing the 95% confidence interval. Coefficient of determination (r^2^) = 0.8514 (**e**) Bar plot showing Pearson correlation coefficients (R) between eccDNA abundance and various genomic instability features. Error bars represent 95% confidence intervals derived from bootstrap analysis. Color coding indicates statistical significance based on confidence interval analysis: blue, red, and grey represent significant positive, significant negative, and non-significant correlations, respectively (95% CI excludes zero). * WGD: whole genome duplication, SVB: structural variant, Cth: chromothripsis-like patterns, CNI: copy number instability per chromosome. **(f)** Box plot showing the enrichment ratio (observed/expected) of eccDNA distribution across different genomic elements. Box plots represent the distribution of enrichment ratios from 1,000 iterations of matched sampling comparisons using 10% subsamples of observed eccDNA with size- and chromosome-matched random regions. Orange boxes indicate normal adjacent tissue (NAT) samples, while blue boxes represent tumor tissue (TT) samples. Error bars show 95% confidence intervals. The red dashed line at 1.0 represents no enrichment (observed equals expected). Values greater than 1.0 indicate enrichment, while values less than 1.0 indicate depletion. **(g)** Metaplots showing normalized signal intensity across eccDNA regions for five chromatin features using size-scaled boundary definitions (15% of eccDNA length, 200bp-1kb range). The x-axis represents relative position from the eccDNA start site (eSS) through the center to the eccDNA end site (eES). Vertical dashed lines demarcate the boundary regions. Shaded areas represent 95% confidence intervals (n=24 samples).

We examined the chromosomal distribution of eccDNA across the genome. Samples with no detectable eccDNA were excluded from this analysis. eccDNA mapped to all chromosomal regions, with notable enrichment in chromosomes 7, 8, 13, and 20 (**Figure 4b and Supplementary Figure 3b**). Positional enrichment analysis (per MB per chromosome) using chi-square goodness-of-fit testing revealed significant regional specificity in eccDNA formation, though enrichment patterns varied among samples (Supplementary Table S4). Ten samples (CRCtx021, 025, 027, 028, 029, 032, 035, 037, 047, 054) showed significant enrichment across all four chromosomes. Eight samples were enriched in three of the four chromosomes (CRCtx024, 030, 033, 034, 036, 039, 040, and 053) and four (CRCtx031, 046, 050, 055) in two. Only one sample (CRCtx023) showed enrichment restricted to chromosome 8, while another (CRCtx022) exhibited no significant over-representation in any of the four chromosomes, displaying a distribution consistent with random genomic origin.

In summary, 42% of samples exhibited enrichment in all four chromosomes, 33% in three, 17% in two, and 8% showed either single-chromosome or no significant enrichment.

We examined the relationship between eccDNA abundance and chromosomal copy number alterations using WGS data from 12 samples. Regression analysis between the number of eccDNA per chromosome per Mb and the average copy number per chromosome revealed a strong positive correlation (**Figure 4c**). This relationship was particularly evident for chromosomes 7, 8, 13, and 20, which showed significant enrichment of eccDNA across multiple samples the same chromosomes identified in our regional bias analysis.

In sample CRCtx021, elevated eccDNA per Mb was observed in chromosomes 7, 8, 13, and 20, directly aligning with corresponding copy number gains (R² = 0.852) (**Figure 4d**). This trend was maintained across all 12 samples (combined R² = 0.714, p < 0.001, 95% CI: [0.672, 0.752]; meta-analysis of 12 samples using Stouffer’s method, **Supplementary Figure 4a**).

We analyzed three instability metrics across the 12 samples: whole genome duplication (WGD), structural variant burden, and chromothripsis-like patterns to assess correlations with eccDNA abundance.

eccDNA abundance showed variable associations with global instability metrics (Figure 2c). WGD events showed no correlation (R = 0.129, 95% CI: [-0.554, 0.645], p = 0.705). Structural variant burden showed a moderate positive correlation (R = 0.406, 95% CI: [0.021, 0.824], p = 0.215), and chromothripsis-like patterns similarly correlated positively (R = 0.425, 95% CI: [0.144, 0.732], p = 0.192). Copy number instability per chromosome showed a moderate negative correlation (R = -0.404, 95% CI: [-0.732, -0.105], p = 0.217).

We analyzed publicly available ATAC-seq and ChIP-seq data for six histone modifications and three transcription regulators from transverse colon samples (ENCODE), using size-scaled boundary definitions (15% of eccDNA length, 200bp-1kb range) and matched random controls (**Supplementary table S5**).

Five chromatin features showed significant enrichment at eccDNA boundaries (FDR < 0.05). ATAC-seq demonstrated the strongest enrichment (1.057-fold, p = 2.27 × 10⁻⁷, Cohen’s d = 2.31). CTCF showed robust boundary enrichment (1.046-fold, p = 2.27 × 10⁻⁷, Cohen’s d = 2.09). Active transcription markers were also enriched: POLR2A (1.031-fold, p = 2.27 × 10⁻⁷, Cohen’s d = 3.60), H3K4me3 (1.021-fold, p = 2.27 × 10⁻⁷, Cohen’s d = 2.88), and H3K27ac (1.017-fold, p = 1.60 × 10⁻³, Cohen’s d = 0.72). Repressive marks (H3K9me3, H3K27me3), H3K36me3, and additional regulatory marks (H3K4me1, EP300) showed no significant enrichment.

Metaplots using 500bp windows confirmed these patterns (**Figure 4g**). Size stratification into small (1-5kb), medium (5-20kb), and large (>20kb) eccDNA revealed distinct chromatin requirements (**Supplementary Figure 5a-d**). Small eccDNA showed dramatically stronger dependencies, with 6-fold higher ATAC-seq signals and pronounced enrichment of all five significant marks. Medium and large eccDNA displayed progressively broader, less feature-specific patterns.

These analyses reveal that eccDNA exhibits nucleosome periodicity, preferential enrichment on chromosomes 7, 8, 13, and 20, and strong correlation with chromosomal amplifications. eccDNA boundaries are enriched for accessible chromatin, CTCF binding sites, and active transcriptional markers, with smaller circles showing stronger chromatin feature dependencies.

## Discussion

In this study, we demonstrate that CRC tumors carry large numbers of eccDNA with overexpressed recurrent oncogenes. We find that one such gene, CXCL5, confers tumor phenotypes when expressed from circular elements in a CRC cancer cell line, and we show that patients with the highest level of eccDNA in their tumors experience relapse. This suggests that eccDNA can provide an evolutionary advantage to tumor cells by amplifying oncogenes or other critical genes. Many eccDNA-borne genes in the TT samples showed higher RNA expression levels compared to those in NAT samples, supporting the hypothesis that eccDNA can give rise to increased gene activity in cancer cells. It has been described before how oncogenes on the large ecDNA are correlated to higher RNA expression and tumor development (7). However, the evidence for eccDNA <100,000 bp’s role in tumorigenesis is limited and indirect. Henssen and coworkers have shown that genes on eccDNA are common and frequently overexpressed in neuroblastoma (3), and Ye and coworkers recently found a similar connection for eight liver tumors (35). To prove that the transcripts derived from the allele on eccDNA and not the chromosomal allele, Henssen, and his team haplotyped transcripts from eccDNA-borne genes and found a strong bias toward the eccDNA-borne alleles. Still, proof that expression from genes on eccDNA is responsible for tumor phenotypes has been lacking (10, 39). We reconstituted one of the eccDNA-bornegenes with the highest increase in RNA expression level in a CRC cell line to test for this. One of the tumor tissue samples exhibited a significantly increased RNA expression level of the chemokine CXCL5 and elevated gene copy numbers on eccDNA compared to the NAT. CXCL5 is overexpressed in many tumors, including colorectal cancer, where it has been implicated in promoting tumor growth, metastasis, and angiogenesis (28, 29, 40). Cells with circular CXCL5 DNA exhibited higher RNA expression levels and demonstrated the ability to affect neutrophils in a cell migration assay. Our findings support the hypothesis that gene amplifications on eccDNA can directly affect tumorigenesis by driving oncogene expression and altering tumor cell phenotypes.

Despite these findings, the presence of genes on eccDNA does not universally correlate with elevated gene expression. A substantial proportion of highly expressed genes in our study were not located on eccDNA, and conversely, many genes found on eccDNA did not exhibit high expression levels. This implies that eccDNA is not the sole determinant of gene expression levels. However, when comparing RNA expression for genes found on eccDNA and ecDNA with the chromosomal linear amplifications, we observed significantly higher RNA expression for genes located on circle amplifications. This suggests that eccDNA provides a more effective platform for selective upregulation of certain genes compared to linear chromosomal amplifications. The effectiveness of circular elements to upregulate gene expression has been shown for the large ecDNA (3, 5, 7, 12), and the ability has been ascribed to missegregation of ecDNA in mitosis and altered chromatinization (7).

A hallmark of genomic instability in cancers, including CRC, is aneuploidy, which is frequently observed in solid tumors (reviewed in (41–44)). In our study, we found that the distribution of eccDNA origins in TT samples was skewed toward specific chromosomes, particularly chromosomes 7, 8, 13, and 20. Interestingly, our WGS analysis revealed amplifications and anomalies (partial aneuploidy) in these same chromosomes. Amplifications in chromosomes 7, 8, 13, and 20 are well-established in CRC (45–50), while the loss or partial loss of chromosomes 8, 17, and 18 is also linked to CRC progression (47). Our study revealed that a substantial portion of eccDNA originates from enhancers and 5’ UTRs, while fewer fragments are derived from exons and introns. This contrasts with a previous CRC study by Chen et al. (2022), which found a higher prevalence of eccDNA originating from exons and introns. Such variations across studies and cancer types reflect the diverse genomic landscapes from which eccDNA can emerge (33, 35, 51, 52). The differences in eccDNA origin may be influenced by the specific tissue types or the stage of cancer progression, highlighting the complexity of eccDNA biogenesis across cancers. The distinct enrichment of CTCF at eccDNA boundaries, along with the presence of open chromatin marks such as H3K4me3 and H3K27ac, suggests that chromatin organization may play a crucial role in the formation or stabilization of eccDNA. CTCF is well-known for its role in establishing higher-order chromatin structure and regulating genomic architecture, often acting as a boundary element that demarcates chromatin domains (53–55). We hypothesize that CTCF binding sites may serve as preferred breakpoints during DNA damage events that generate eccDNA, similar to their role in translocation breakpoints (54). The enrichment of the active chromatin marks H3K4me3 and H3K27ac, which are typically associated with active enhancers and promoters, further supports the idea that eccDNA formation is biased toward regions of open and transcriptionally active chromatin (56, 57). These regions are often hotspots for regulatory activity (58, 59), suggesting that active chromatin features may predispose certain regions of the genome to eccDNA production (60, 61).

Our data also revealed an increased amount of eccDNA in all tested CRC tumor samples compared to non-tumorous samples, aligning with previous studies on colon and other tumors (33, 62, 63). The presence of eccDNA, particularly the ecDNA subpopulation, is linked to DNA damage, oncogene amplification, and cancer heterogeneity (5, 15, 64–67) and has also been documented in the CRC (32, 33, 68). Furthermore, smaller eccDNA has been associated with DNA damage and genome instability (69). Notably, we observed that a higher amount of eccDNA correlates with poor relapse-free survival in patients. In earlier works, cancer patients detected with the large ecDNA carrying oncogenes or having complex ecDNA had a lower 5-year survival in comparison with patients without the ecDNA (3, 12, 14, 70, 71). It has been shown that ecDNA serves as a fast adaptation tool for better survival for cancer cells, which leads to poorer survival for cancer patients (5). Our findings show that not only the big complex ecDNA molecules but also the broad-size (<100,000 bp) spectrum eccDNA are linked to tumor survival and development.

### Broader Biological Significance and Future directions

Our findings demonstrate that eccDNA, including smaller circular DNA elements (<100,000 bp), plays a more significant role in cancer biology than previously recognized. We show that eccDNA can serve as a platform for oncogene amplification and expression in CRC, with direct implications for tumor phenotypes, as demonstrated through our CXCL5 functional studies. The preferential formation of eccDNA from active chromatin regions and its correlation with chromosomal copy number variations suggests a complex interplay between genomic instability, chromatin state, and eccDNA biogenesis. Importantly, the correlation between eccDNA abundance and poor patient outcomes indicates that eccDNA may serve as both a prognostic marker and a potential therapeutic target. Understanding the mechanisms of eccDNA formation and its contribution to tumor evolution could lead to novel therapeutic strategies aimed at preventing or reducing eccDNA-mediated gene amplification in cancer cells. Future studies should focus on developing methods to specifically target eccDNA-bearing cells or prevent eccDNA formation, potentially offering new approaches to combat cancer progression and drug resistance.

### Limitations and Methodological Considerations

The cross-sectional design precludes causal inferences about amplification-eccDNA relationships. The small cohort size limits statistical power for clinical associations and generalizability across populations. The survival trend requires validation in larger, multi-institutional studies. Technical challenges in detecting small eccDNA may influence observations, though validation experiments support biological relevance. While our findings are consistent with chromatin-guided eccDNA formation, alternative models including amplification-driven eccDNA excision cannot be excluded based on available data.

In conclusion, eccDNA represents a prevalent and functionally significant feature of colorectal cancer with distinctive chromatin associations and gene amplification capabilities across all size ranges. These findings provide a foundation for future mechanistic studies and potential therapeutic strategies.

## Materials and Methods

### Sample collection and description

Our study cohort comprises 25 samples of paired colorectal tumor tissue (TT) and adjacent normal adjacent tissue (NAT) samples (N =50). The average age of the cohort sample was 66 years; with 14 females and 11 males with varying tumor stages (2a, 2b, 3a, 3b). The characteristics of the cohort are summarized in **Supplementary Table S1**.

This study was performed in accordance with the declaration of Helsinki. Patient tissue samples were obtained from the REBECCA biobank in Denmark and all samples were stored at -80 °C. The REBECCA study protocol was approved by the Ethics Committee of the Capital Region of Denmark (VEK j.nr. H-2-2013-078) and the Danish Data Protection Agency (j. nr. 2007-58-0015, HEH-014-044, I-Suite nr. 02771). Patients were recruited following ethical guidelines, and informed consent was obtained before sample collection. To ensure privacy, all samples underwent anonymization procedures before experimental processing. Anonymization involves assigning unique identifiers to each sample that are unlinkable to personal information.

### Cell culture

The human SW620 colon cancer cell line and THP-1 monocyte cell line (mycoplasma tested) were cultured in RPMI 1640 Medium (ThermoFisher, USA), enriched with 10% Fetal Bovine Serum (ThermoFisher, USA) and 1% penicillin-streptomycin (ThermoFisher, USA), and incubated in a humidified incubator at 37 °C with 5% CO_2_. Cells were cultured to 70–90% confluence in 10 cm² Petri plates and passaged every 2-3 days to maintain this cell density.

### Plasmids and linear DNA quality control

For the quality control testing, we spiked-in a control mixture consisting of plasmids ((50,000 copies of p4339 (5064 bp), 10,000 copies of pBR322 (4,361 bp) (NEB, USA)). All plasmids were maintained in Escherichia coli and purified with a standard plasmid midi-prep kit (NucleoBond® Xtra Midi, MACHEREY-NAGEL, DE). For quality control assessment, we used the standard protocol as per the manufacturer of qPCR assay with SYBR Green master mix (2x) (Applied Biosystems, USA) with primers from **Supplementary Table S6** and the results are shown in **Supplementary Figure 5e**.

### Total DNA extraction, mitochondrial DNA and linear DNA removal from tissue samples

DNA from all tissue samples (6 mg) were extracted with MagAttract HMW DNA Kit (Qiagen, DE) as described in the manufacturer’s protocol. Total DNA from all samples was subjected to treatment with a plasmid-safe Exonuclease V for 5 days to remove linear DNA. We also removed mtDNA by using CRISPR/Cas9 system as described in Feng et al. 2022 with small changes (Feng et al., 2022). In short, at first, for 3 days prior mtDNA removal, we digested linear DNA by Exonuclease V. 20 μl of 10X NEB buffer 4, 20 μl of 10 mM ATP, 6 μl of Exonuclease V and 4 μl water (NEB, USA) were added per reaction and incubated for 3 days at 37 °C (every day adding extra 1.4 μl of 10X NEB buffer 4, 10 μl of 10 mM ATP and 3 μl of Exonuclease V), at the end heat-inactivated for 30 min at 70 °C). DNA was then cleaned using AMPure XP magnetic beads (Beckman Coulter, USA). mtDNA was linearized by Cas9 Nuclease, *S. pyogenes* kit (NEB, USA) according to the manufacturer’s protocol. All gRNAs were designed as in Feng et al. 2022 (Feng et al., 2022). For CRISPR/Cas9 combining, 100 nM of Cas9 protein and 100 nM of sgRNA were mixed in a 1×NEB 3.1 buffer and incubated at 25 °C for 30 min. Then, sample DNA was added, mixed thoroughly, and incubated at 37 °C for 4 hours. Then, we inactivated the Cas9: sgRNA cleavage system by heating the sample for 10 min at 65 °C. After this step, Exonuclease V digestion was performed again to remove the now linearized mtDNA (2.1 μl of 10X NEB buffer 4, 15 μl of 10 mM ATP and 4.5 μl of Exonuclease V were added every day to the same reaction and incubated for 2 days at 37 °C, then heat-inactivated for 30 min at 70 °C). Followed by DNA cleaning using 1.8x ratio of AMPure XP magnetic beads that were added to each sample, mixed by pipetting 10x and incubated at room temperature for 5 min. The samples were put on a magnetic rack for 3 min, following which the supernatant was discarded. The samples were then washed twice with 200 µL 75-80% ethanol and dried until the beads were slightly moist. The beads were then resuspended in 15 µL of 10 mM Tris-HCl (pH=8) (ThermoFisher, USA) buffer and mixed by pipetting 10x, before being incubated for 5 min at 50 °C. This was followed by a 2 min bead separation step on the magnetic rack, after which 13 µL of the eluate was taken out and transferred to a clean new 1.5 mL DNA Lobind tube (Eppendorf, DE) (2 µL were left behind). The elution step was then repeated using 12 µL elution buffer and combined with the other elute for a final sample volume of 25 µL. For DNA concentration measurements and quality control, 10 µL of each sample was transferred to a new PCR tube.

### Rolling-circle amplification of eccDNA for Circle-Seq for tissue samples

All the purified eccDNA samples (15 µL) were then used as a template for Ф29 polymerase reactions (4BB™ TruePrime® RCA Kit) (4basebio PLC, UK), which was added in accordance with the manufacturer’s instructions and incubated for 48 h at 30 °C.

### RNA extraction from tissue samples

RNA was extracted from the last 25 paired samples. For tissue under 5 mg, we used the miRNeasy Tissue/Cells Advanced Micro Kit (Qiagen, DE), and for tissue up to 30 mg, we used the miRNeasy Tissue/Cells Advanced Mini Kit (Qiagen, DE), as stated in the manufacturer’s protocol. For quality control, we used the Agilent RNA 6000 Pico Kit (Agilent Technologies, USA).

### eccDNA, whole genome DNA, and RNA library preparation and sequencing

For the eccDNA sequencing, a portion of each Ф29-amplified DNA sample was diluted to 15 ng/µL in 10 mM Tris-HCl (pH=8) for a total volume of 100 µL and then sonicated (4 cycles of 20sec/30sec (on/off time) using a Bioruptor (Pico, Diagenode, USA).

For the WGS, total DNA was diluted to 15 ng/µL in 10 mM Tris-HCl (pH=8) for a total volume of 100 µL and then sonicated in the same manner as for plasma samples. The libraries were then prepared using the NEBNext Ultra II DNA Library Prep Kit for Illumina and NEBNext Multiplex Oligos for Illumina (NEB, USA) in accordance with the manufacturer’s protocol. For the sonication step and library quality control, we used the Agilent DNA 1000 Kit (Agilent Technologies, USA).

Following library preparation, all eccDNA samples were multiplexed and sequenced on a Novaseq 6000, S4 flow-cell (Illumina, USA) as 2×150-nucleotide paired-end reads, with an average of 157 million single reads per sample. WGS samples were sequenced with Novaseq 6000, S1 flow-cell as 2×150-nucleotide paired-end reads, with an average of 62 million single reads per sample (all sequencing was done at Rigshospitalet, Denmark).

For RNA-seq, we used NEBNext rRNA Depletion Kit v2 (Human/Mouse/Rat) and NEBNext Ultra II Directional RNA Library Prep Kit for Illumina; for barcoding, we used NEBNext Multiplex Oligos for Illumina (96 Unique Dual Index Primer Pairs) (NEB, USA) as stated in manufacturer’s protocol. RNA was multiplexed and sequenced on a Novaseq 6000, SP flow cell as 2×75-nucleotide paired-end reads, with an average of 25 million single reads per sample (all sequencing was done at Rigshospitalet, Denmark).

### eccDNA copy number variation detection by qPCR

To check for copy number variations for CXCL5 gene with high RNA expression in cancer tissue samples, we used qPCR assay to measure relative copy number in TT and NAT samples by using SYBR™ Green PCR Master Mix (2x) (Thermo Scientific, USA) and a primer pair from **Supplementary Table S6**. For qPCR, after a denaturation step at 95 °C for 10 min, samples were incubated for 40 cycles at 95 °C for 15 s and at 60 °C for 30 s. For eccDNA-purified DNA concentration was unknown as it was too low to detect by Qubit. Serial dilution standard curves for qPCR were used to determine CNV. Relative CN in eccDNA-purified DNA=CN CXCL5 gene/CN of spike-in plasmid. Each sample was run in triplicates. A melting curve analysis was performed to verify the specificity and identity of the qPCR products. Removal of linear DNA was confirmed during the same qPCR runs by ALB gene primers and with an additional PCR reaction with Cox5B primers using DreamTaq PCR Master Mix (2X) (Thermo Scientific, USA) standard protocol with modifications by adding 5% of DMSO and 2 nM of MgCl_2_, and 40 cycles and visualized on 2% agarose gel (SeaKem LE Agarose, Lonza Group, CH) with 60x GelRed Nucleic Acid Gel Stain (Biotium, USA) (**Supplementary Figure 9 c,e**).

### [CXCL5^circle^] eccDNA validation with PCR

To confirm the presence of the [CXCL5^circle^] in the CRCtx028 sample, we performed PCR targeting the junction site of the circle. The PCR was conducted using DBdirect™ PCR Gel Mix SuperSens (2x) (DIANA Biotechnologies, CZ) according to the manufacturer’s instructions, with [CXCL5^circle^] primers (final concentration of 5 nM) listed in **Supplementary Table S6**. We used rolling-circle amplified DNA from the CRC-028 patient’s tumor tissue sample and normal adjacent tissue sample, diluted to a concentration of 45 ng/µL (2.25 ng/µL final concentration). The PCR amplicons were visualized on 1% agarose gel (SeaKem LE Agarose, Lonza Group, CH) with 60x GelRed Nucleic Acid Gel Stain (Biotium, USA).

### Reverse transcription

iScript™ cDNA Synthesis Kit (Bio-Rad, USA) was used for the reverse transcription according to the manufacturer’s guidelines. Briefly, 20 μl reaction system was prepared, including 500 ng of total RNA, 4 μl of 5x iScript reaction mix, 1 μl of iScript reverse transcriptase and nuclease-free water. To initiate the reverse transcription operations, the temperature was first set to 25°C for 5 min, then raised to 46 °C for 20 min, and finally increased to 95 °C for 1 min to inactivate the reaction.

### Droplet Digital PCR

Droplet Digital PCR (ddPCR) reactions were set up using the subsequent methodology in a 20 μl reaction system, including 5 ng of genome DNA, 2 pmol of each prime, 10 μl QX200TM ddPCR^TM^ EvaGreen Supermix (Bio-Rad, USA), and sterile water. The PCR mixture was then put in a Bio-Rad QX200 Droplet Generator (Bio-Rad, USA) to produce droplets. Subsequently, a 40 μl droplet was placed onto a Bio-Rad 96-well PCR plate and sealed with foil, heating it to 170 °C for 4 seconds. Next, 98 °C for 5 min, 40 cycles of 95 °C for 30 s and 60 °C for 1 min, followed by 4 °C for 5 min, 90 °C for 5 min, and a 4 °C hold, were used for the PCR. For every phase, a ramp rate of 2 °C/s was employed. The droplets were then read using the Bio-Rad QX200 ddPCR apparatus. The data visualization and analysis software utilized was Bio-Rad QuantaSoftTM Analysis (Version 1.0.596).

### Nucleofection

CXCL5 circular DNA was synthesized by GenScript Europe company (Rijswijk, NL), and delivered to SW620 cells using the SE Cell Line 4D-Nucleofector^TM^ X Kit S (Lonza Group, CH) in accordance with the manufacturer’s instructions. Briefly, the SW620 cells were trypsinized, quenched with culture medium (1% penicillin-streptomycin and 10% FBS), and 10^6^ cells were counted and centrifuged at 500 rpm for 5 min. After a single PBS buffer wash, the cells were resuspended in 20 μl of the electroporation solution, and 1 μg of CXCL5 circular DNA was added. Then, the cell suspension was moved to a nucleocuvette and electroporated in a Lonza 4D-Nucleofector device under the CM130 program. Subsequently, the cells were seeded in a six-well plate with pre-warmed culture media and underwent two days of culturing.

### Boyden Chamber cell migration assay

Two days post-nucleofection, the culture medium of the SW620 cells was transferred to the bottom chamber of a 24 mm Transwell® with an 8.0 µm pore polycarbonate membrane insert (Corning, USA). Simultaneously, 2×10^6^ THP-1 cells were seeded in the upper chamber of the same Transwell® for a 6-hour migration. After the migration period, the upper chambers were removed, and the cell numbers in the lower chambers were counted.

### RNA expression analysis

The quality of the RNA-sequencing data was assessed using Fastqc (RRID: SCR_014583) v0.11.9 (https://www.bioinformatics.babraham.ac.uk/projects/fastqc/), following which the low-quality reads and adapters were trimmed using fastp (RRID: SCR_016962) v0.20.1 (https://github.com/OpenGene/fastp). The quantification of the transcripts was obtained using Kallisto (RRID: SCR_016582) v0.50.1 (https://pachterlab.github.io/kallisto/) as Transcripts per Kilobase Million (TPM) based on the GRCh38 reference genome. DEseq2 was used to calculate the fold change for each tumor tissue sample in comparison to the collectively grouped normal adjacent tissue samples.

### Gene set enrichment analysis

500 differentially expressed genes (DEGs) from DEseq2 were used for the gene set enrichment analysis (GSEA). Two different datasets of the MSigDB resources (http://www.gsea-msigdb.org/gsea/msigdb/index.jsp) Hallmark and C6 (oncogenic gene sets) were used through the GSEA (RRID: SCR_003199). Gene sets with an FDR-corrected p <0.01 were considered significantly enriched, and the results were plotted using custom Python scripts.

### eccDNA identification

We employed a mapping-based approach to identify chromosomal-derived eccDNAs, following the pipeline described by Feng et al. (2022). An event was considered a circle if supported by at least two sequence reads indicating the presence of a chimeric alignment, discordantly paired-end mappings, or soft-clipped reads spanning the circle breakpoint. Notably, recurrent eccDNAs such as mitochondrial DNA were counted as single events. The identified eccDNAs were assigned further confidence levels based on coverage criteria: PASS1: At least 95% of positions within the putative eccDNA circle were covered by reads. PASS2: Mean coverage within the circle was at least twice the mean coverage in neighboring regions in addition to fulfilling the PASS1 criteria.

Gene annotations were based on the GRCh38 assembly, obtained from ENSEMBL (http://ftp.ensembl.org/pub/release-110/gtf/homo_sapiens/Homo_sapiens.GRCh38.110.gtf.gz). We used BedTools v2.30.0 (RRID: SCR_006646) intersect to determine gene containment within eccDNAs. An eccDNA was considered to contain a whole gene if the gene’s start position was greater than the eccDNA’s start position and the gene’s end position was less than the eccDNA’s end position.

### WGS analysis

Raw sequencing reads were assessed for quality using FastQC (RRID: SCR_014583) v0.11.9. Adapter sequences and low-quality bases (Phred score < 30) were trimmed using Trimmomatic (RRID: SCR_011848) v0.38. Clean reads were aligned to the human reference genome (GRCh38) using BWA-MEM (RRID: SCR_010910) v2.2.1 with default parameters. The resulting SAM files were converted to BAM format, sorted, and indexed using SAMtools (RRID: SCR_002105) v1.18. PCR and optical duplicates were marked using Picard MarkDuplicates (RRID: SCR_006525) v2.9.1.

### Copy Number Variation Analysis

Copy number analysis was performed using CNVkit (SCR_021917) v0.9.9. We used duplicate-removed BAM files as input, with the hg38 human reference genome as the baseline. To improve accuracy and reduce false positives, we applied an access list to exclude problematic regions of the genome based on a modified version of the ENCODE blacklist regions for hg38. Gene-level annotations were incorporated using RefSeq gene definitions (refFlat.txt) to provide biological context to the detected CNVs. Custom Python scripts were used for further downstream analysis and visualization.

### Whole Genome Doubling Detection

WGD events were identified using a multi-criteria approach based on CNVkit-derived copy number profiles. For each sample, we calculated the fraction of the genome showing copy number gains (log2 ratio > 0.58, corresponding to copy number ≥ 1.5) by summing the length of gained segments divided by total analyzed genome length. WGD status was assigned when ≥50% of the genome showed coordinated copy number increases and ≥10 chromosomes displayed gains in >30% of their length. A quantitative WGD score was calculated as the genome-wide fraction of gained segments to enable correlation analysis with eccDNA abundance.

### Structural Variant Burden Quantification

Structural variant burden was estimated from copy number breakpoint density using CNVkit segmentation data. Breakpoints were identified as positions where consecutive segments on the same chromosome showed copy number changes exceeding 0.3 in log2 space. For each sample, we calculated: (1) total estimated breakpoints across all chromosomes, (2) copy number variance as a measure of segmental instability, and (3) per-chromosome copy number instability defined as the variance of log2 ratios within each chromosome. A composite SV burden score was computed by combining breakpoint density and copy number variance (breakpoints + variance × 100) to provide a single metric for correlation analysis.

### Chromothripsis-like Event Detection

Chromothripsis-like events were identified by analyzing patterns of clustered copy number oscillations within individual chromosomes. For each chromosome, we calculated the frequency of high-amplitude copy number changes (|Δlog2| > 0.5) between consecutive segments. Chromosomes were classified as chromothripsis-like when >30% of segments showed high-amplitude oscillations, indicating extensive local rearrangement. A chromothripsis score was calculated as the proportion of chromosomes per sample showing chromothripsis-like patterns, enabling quantitative correlation with eccDNA abundance.

### Genomic Instability Profile Integration

To assess combined genomic instability effects, we generated a composite instability score by standardizing and summing WGD scores, SV burden scores (normalized by dividing by 100), and chromothripsis scores. This combined metric was correlated with total eccDNA abundance to test whether multiple instability mechanisms collectively influence eccDNA formation patterns.

Correlation confidence intervals were calculated using bootstrap resampling (10,000 iterations) to account for small sample size effects and provide robust estimates of statistical significance. Bootstrap confidence intervals were prioritized over Fisher-z transformation due to their superior performance with small samples. Correlations were considered statistically significant if the 95% bootstrap confidence interval excluded zero.

### Amplicon Architect

Regions exhibiting copy numbers greater than five were selected as putative intervals for ecDNA reconstruction using Amplicon Architect (v1.2). Amplicon Architect was used to assemble focal amplifications, identifying and characterizing the structure of amplified genomic regions. The resulting assemblies were further classified using Amplicon Classifier (v0.4.11) (Luebeck et al., 2023), which was run with default parameters to categorize the amplicon structures.

### eccDNA Chromosomal Distribution

To determine the genomic distribution of eccDNA across chromosomes, we first normalized the number of eccDNA occurrences per chromosome to per Mb, followed by z-score normalization to account for differences in chromosome length. To assess whether the occurrence of eccDNA in each chromosome was significantly higher than expected by chance, a chi-square goodness-of-fit test was performed, comparing observed eccDNA distributions to expected distributions based on chromosome length.

### Copy Number Variation Analysis

CNV data were processed from CNVKit results in BED file format. CNV per Mb was calculated by binning the hg38 reference genome using the bedtools makewindows function, followed by intersecting the log2 ratio values from the CNV BED files with genomic bins. This approach provided normalized CNV measurements across all chromosomes for correlation analysis with eccDNA density.

### Correlation and Regression Analysis

Linear regression analysis was performed to quantify relationships between eccDNA density per chromosome and corresponding genomic metrics (copy number alterations and gene density) using the linregress function from scipy.stats. Pearson correlation coefficients were calculated to assess the strength of linear relationships, with R² values determined to quantify the proportion of variance explained by each model.

### Meta-Analysis

For multi-sample analysis, individual correlation results were combined using Stouffer’s method for p-value combination and weighted averaging for R² values. Individual p-values were converted to Z-scores and combined using sample size weights (wi = ni - 2, representing degrees of freedom for linear regression): Zcombined = (Σ wi × Zi) / √(Σ wi). The combined R² was calculated as a weighted average: R²combined = (Σ wi × R²i) / (Σ wi). Confidence intervals for combined effect sizes were calculated using Fisher’s Z transformation with appropriate back-transformation to the correlation scale. For meta-analysis results, combined p-values were calculated using Stouffer’s method, and 95% confidence intervals were reported for all combined effect sizes.

### Genomic Instability Metrics

Genomic instability was quantified using four metrics: (1) whole genome duplication fraction (proportion of genome with copy number gains), (2) structural variant burden (breakpoint density and altered genome fraction), (3) chromothripsis-like patterns (oscillating copy number changes), and (4) copy number instability (variance of log2 ratios per chromosome).

Pearson correlations between eccDNA abundance and instability metrics were calculated for 11 samples with complete data. Given the small sample size, 95% confidence intervals were estimated using bootstrap resampling (10,000 iterations) rather than parametric methods. Correlations were considered significant if bootstrap confidence intervals excluded zero. Bootstrap methods provide robust significance testing for small-sample correlations without distributional assumptions.

### Chromatin mark enrichment

Publicly available ATAC-seq and ChIP-seq data generated from human transverse colon tissue were downloaded from ENCODE as BigWig and Bed files. Accession numbers for the datasets used are provided in **Supplementary Table S5**

### eccDNA Filtering and Size Stratification

eccDNA regions were filtered to include only those ≥1kb in length to ensure reliable boundary analysis. For size stratification analysis, eccDNAs were classified into three categories: small (1-5kb), medium (5-20kb), and large (>20kb). This stratification was performed to investigate potential size-dependent chromatin formation mechanisms.

### Size-Scaled Boundary Definitions

To address the biological heterogeneity of eccDNA sizes, we implemented adaptive boundary definitions rather than fixed-size regions. Boundary regions were defined as 15% of each eccDNA’s length, with minimum and maximum limits of 200bp and 1kb, respectively. This approach ensures biologically relevant boundary analysis across the diverse eccDNA size spectrum while maintaining statistical power. Internal regions were defined as the middle 50% of each eccDNA, excluding boundary regions to prevent overlap.

### Matched Random Controls

Size-matched random genomic controls were generated to validate the specificity of chromatin enrichments. For each eccDNA in each sample, we generated three random genomic intervals of identical length, ensuring a minimum 5kb distance from any eccDNA to avoid confounding effects. Random intervals were distributed genome-wide proportional to chromosome sizes, creating a robust null distribution for comparison.

### Signal Quantification and Normalization

Chromatin signals were extracted from BigWig files using pyBigWig, with mean signal values calculated for each defined region (boundary, internal, external). To account for local chromatin context differences, signals were normalized relative to ±10kb flanking windows around each eccDNA. Fold enrichment was calculated as the ratio of boundary signal to internal signal for each eccDNA.

### Statistical Analysis

Boundary enrichment was assessed using the Wilcoxon signed-rank test, comparing fold enrichment values against the null hypothesis of no enrichment (fold change = 1). Effect sizes were calculated using Cohen’s d approximation. To control for multiple testing across ten chromatin features, we applied the Benjamini-Hochberg false discovery rate (FDR) correction with α = 0.05. Statistical analyses were performed in Python using scipy.stats.

### Metaplot Generation

To visualize spatial chromatin patterns, we generated metaplots showing normalized signal intensity across eccDNA regions. Each eccDNA was divided into 100 bins with additional 20-bin flanking regions (500bp upstream/downstream). Profiles were aggregated across all samples and smoothed using Gaussian filtering (σ = 1.0) for visualization. Size-stratified metaplots were generated separately for each eccDNA size category.

### Genomic annotation of eccDNA regions

To assess whether eccDNA formation occurs preferentially in specific genomic features, we performed matched sampling analysis comparing observed eccDNAs with size- and chromosome-matched random genomic regions. We randomly sampled 10% of eccDNAs from each sample (or 50,000 regions for samples with >500,000 eccDNAs) and generated an equal number of random regions matching the exact chromosome and size distribution of the sampled eccDNAs. This process was repeated for 1,000 iterations to generate robust statistics. Genomic annotations for hg38 were obtained using the annotatr package v3.19 (10.18129/B9.bioc.annotatr), including promoters (1-5kb upstream of transcription start sites), 5’ UTRs, 3’ UTRs, exons, introns, intergenic regions, and FANTOM5 permissive enhancers. For each iteration, we calculated the number of eccDNAs and matched random regions overlapping each genomic feature. Enrichment was calculated as the ratio of observed to expected overlap frequencies, normalized by feature size (regions per Mb). Statistical significance was assessed using the empirical distribution of enrichment values across iterations, with p-values calculated as the proportion of iterations where enrichment deviated from 1.0. Multiple testing correction was performed using the Benjamini-Hochberg method. To examine size-dependent patterns, we stratified eccDNAs into small (<1 kb) and large (>10 kb) categories and repeated the analysis. All analyses were performed separately for tumor and normal samples to identify tissue-specific patterns. Confidence intervals (95%) were calculated from the distribution of enrichment values across iterations.

### Clinical features and survival analysis

Clinical characteristics of patients, including age, sex, cancer stage, and tumor location, were compared to the number of eccDNAs present in tumor tissues using a linear regression model. Patients were further stratified into eccDNA-high and eccDNA-low, based on a cutoff of 5 × 10⁶ eccDNAs, representing the average eccDNA number across the cohort. For survival analysis, patients were stratified into low and high eccDNA groups based on the median tumor eccDNA count. Recurrence-free survival was defined as the time from surgery to first documented disease recurrence or last follow-up for patients without recurrence. Survival curves were constructed using the Kaplan-Meier method and compared between groups using the log-rank test. Cox proportional hazards regression was initially attempted to calculate hazard ratios; however, due to complete separation in the data (zero events in the low eccDNA group), hazard ratio estimation was not feasible. Model discrimination was evaluated using Harrell’s concordance index (C-index) with 95% confidence intervals calculated using bootstrap methods. R packages survival (v.3.4.0) and survminer (v.0.4.9) were used for the analysis.

### Random Dataset Generation and Permutation Analysis

To generate random datasets for comparative analysis with the eccDNA regions, we utilized the read_regions and randomize_regions functions. Using the randomize_regions function, we generated 25 random datasets for each sample with random genomic regions, allowing for overlaps and maintaining the per-chromosome distribution of the original eccDNA regions. The number of eccDNA-borne genes was computed for each random dataset using bedtools intersect. The difference in means between the observed tumor data and the randomly generated datasets was employed as the test statistic for evaluating enrichment.

The NumPy.random.shuffle in Python function was utilized to shuffle the combined dataset (which included both the tumor and random data) and calculate the difference in means for the permuted groups. This shuffling process helps assess how the observed differences could arise under the null hypothesis that no true effect exists. We executed 10,000 permutations to generate a distribution of differences in means, which serves as a reference to evaluate the observed test statistics. The p-value was calculated by determining the proportion of permuted differences as extreme as or more extreme than the observed difference in means. A lower p-value indicates stronger evidence against the null hypothesis, suggesting significant enrichment of eccDNA-borne genes in the tumor samples compared to the random datasets.

### Gene Expression Analysis

We identified genes within eccDNA from Circle-seq data and linearly amplified genes from Amplicon Architect analysis of WGS data across 12 tumor samples. Genes were classified into three groups: eccDNA-borne, linearly amplified, and non-amplified (control). RNA expression data were Z-score normalized for all identified genes across corresponding samples. To evaluate differences in gene expression between groups, a Kruskal-Wallis test was performed using the scipy.stats module.

### Functional Annotation of Differentially Expressed Genes

To determine the functional relevance of the differentially expressed genes, we analyzed the subset of DEGs for their involvement in cancer-related functions. A comprehensive list of 2,769 cancer-related genes was obtained from four online sources: Network of Cancer Genes and Healthy Drivers (http://ncg.kcl.ac.uk/), Cancer Genetic Web (https://www.cancer-genetics.org/), Oncogene database (https://ongene.bioinfo-minzhao.org/browse_gene.html), and Federal Office for Consumer Protection and Food Safety https://zag.bvl.bund.de/onkogene/index.jsf?dswid=8028&dsrid=703. A binomial test was employed using scipy.stats.binomtest function from Python to evaluate whether the number of cancer-related DEGs within the eccDNA dataset was significantly higher than expected by chance.

### Statistical analysis

All statistical analyses were conducted using Python 3 or R version 4.1.2. Continuous variables were compared between tumor and normal tissues using the Wilcoxon rank-sum test due to non-normal distribution of eccDNA counts. Effect sizes were calculated using Cohen’s d. Correlations between eccDNA levels and clinical characteristics were assessed using linear regression models using the following formula:

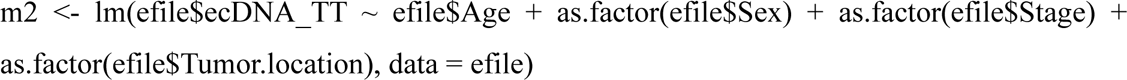

### Statistical Significance

Statistical significance was set at p < 0.05 for all analyses. All tests were two-sided unless otherwise specified. Results are presented as mean ± standard deviation or median (interquartile range) as appropriate. The statistical significance of differences between [CXCL5^circle^] experimental groups was assessed by one-way ANOVA and subsequent Tukey’s HSD (honestly significant difference) test using the python scipy library.

### Data Availability

The datasets supporting the conclusions of this article are available in the Sequence Read Archive (SRA) under BioProject accession number PRJNA1186580 (https://www.ncbi.nlm.nih.gov/bioproject/PRJNA1186580). The codes used to generate the results are available at https://github.com/users/judithhariprakash/projects/.

## Author Contributions

**Judith Mary Hariprakash**: Conceptualization (equal); software (equal); investigation (lead); data curation (lead); writing – original draft (lead); visualization (lead). **Egija Zole**: Conceptualization (equal); investigation (equal); writing – original draft (equal); **Weijia Feng**: Investigation (equal); **Dan Hao**: Software (equal); data curation (equal); **Lasse Bøllehuus Hansen**: Software (equal); data curation (equal); **Nirmalya Bandyopadhyay**: Software (equal); data curation (equal); **Marghoob Mohiyuddin**: Software (equal); data curation (equal); **Astrid Zedlitz Johansen**: Resources (equal); **Julia Sidenius Johansen**: Resources (equal); funding acquisition (equal); **Birgitte Regenberg**: Conceptualization (lead); writing – original draft (lead); visualization (equal); supervision (lead), funding acquisition (lead); **All authors:** data interpretation, critical revision of the manuscript and final approval of the submitted version.

## Acknowledgments

We thank the support from the members of the Regenberg lab, especially the technical assistance from Sefa Alizadeh. This work has been performed using Computerome and computing resources at the core facility for Biocomputing at the Department of Biology, University of Copenhagen.

## Supplementary Figure legends

**Supplementary Figure 1.**
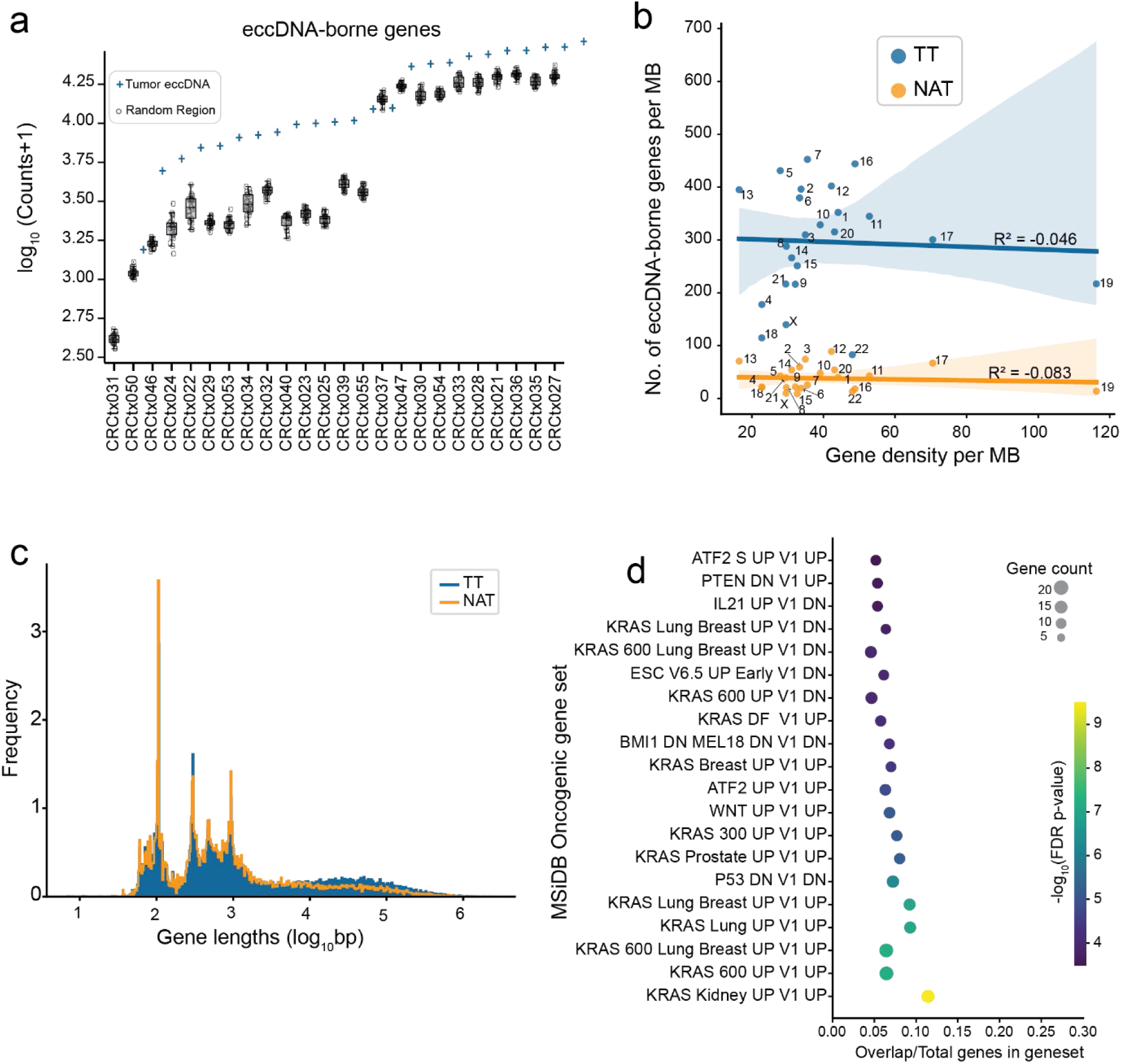
Characterization of eccDNA-borne genes. (**a**) Comparison of eccDNA-borne genes and random genomic regions across patient samples. The y-axis shows log_10_(counts+1) of eccDNA, while the x-axis represents different TT samples. Blue plus sign indicates the number of eccDNA-borne genes in TT sample. The box plot shows the distribution of the number of eccDNA-borne genes in 25 synthetic datasets for each sample. Each box represents the interquartile range (IQR) containing the middle 50% of the data, with the horizontal line inside the box indicating the median. The whiskers extend to the minimum and maximum values within 1.5 times the IQR. Individual data points are plotted as dots, representing specific samples. **(b)** Regression plot showing the no. of eccDNA-borne genes per MB to the gene density per MB. Each point represents an individual chromosome. Blue and yellow dots represent TT and NAT samples respectively. The blue and yellow line indicates the linear regression fit, with the shaded area showing the 95% confidence interval. Coefficient of determination (r^2^) for each category is mentioned in the figure. **(c)** Distribution of gene lengths (log_10_ base pairs) for eccDNA-borne genes in TT (blue) and NAT (yellow) samples. The x-axis represents gene length, and the y-axis shows frequency. (**d**) The scatter plot shows the over-representation of differentially expressed eccDNA-borne genes in the MSigDB Oncogenic gene set. The x-axis represents the ratio of the overlapping genes to the total number of genes in the pathway. The size of the circle denotes the number of genes in overlap, and the color shows the negative logarithmic adjusted p.

**Supplementary Figure 2.**
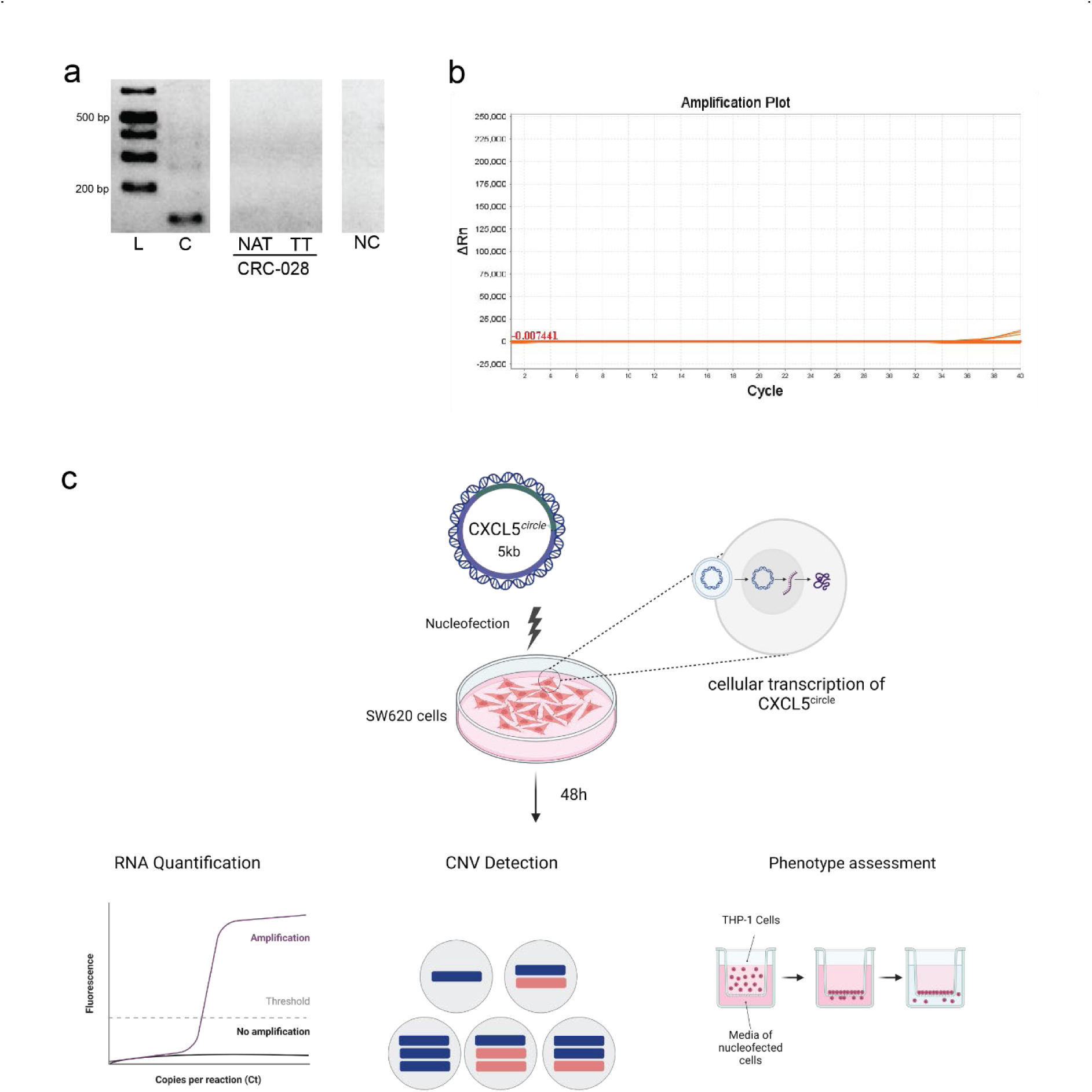
Experimental validation. **(a)** An agarose gel electrophoresis of PCR amplification products was performed to verify the presence of linear DNA following the linear DNA removal process. L, 1 kb plus ladder; TT, tumor samples; NAT, non-tumorous samples; PC, positive control; NC, negative control; bp, base pairs. **(b)** qPCR amplification of the presence of linear DNA following the linear DNA removal process. The ΔRn value represents the difference between the Rn value of the experimental reaction and the Rn value of the baseline signal produced by the instrument. Rn, normalized reporter signal. **(c)** Experimental workflow for studying the effects of [CXCL5*^circle^*] in SW620 cells: Nucleofection: The top of the figure shows a representation of the CXCL5 synthetic construct with 3,035 bp long gene with its 1,845 bp promoter being introduced into SW620 cells via nucleofection. Incubation: The cells are then incubated for 48 hours. Analysis: The workflow branches into three types of analyses: RNA Quantification, CNV Detection and Phenotype assessment using Boyden Chamber cell migration assay in which monocytes (THP1 cells) exposed to medium from SW620 with and without [CXCL5*^circle^*] to test their ability to migrate through a polycarbonate membrane.

**Supplementary Figure 3.**
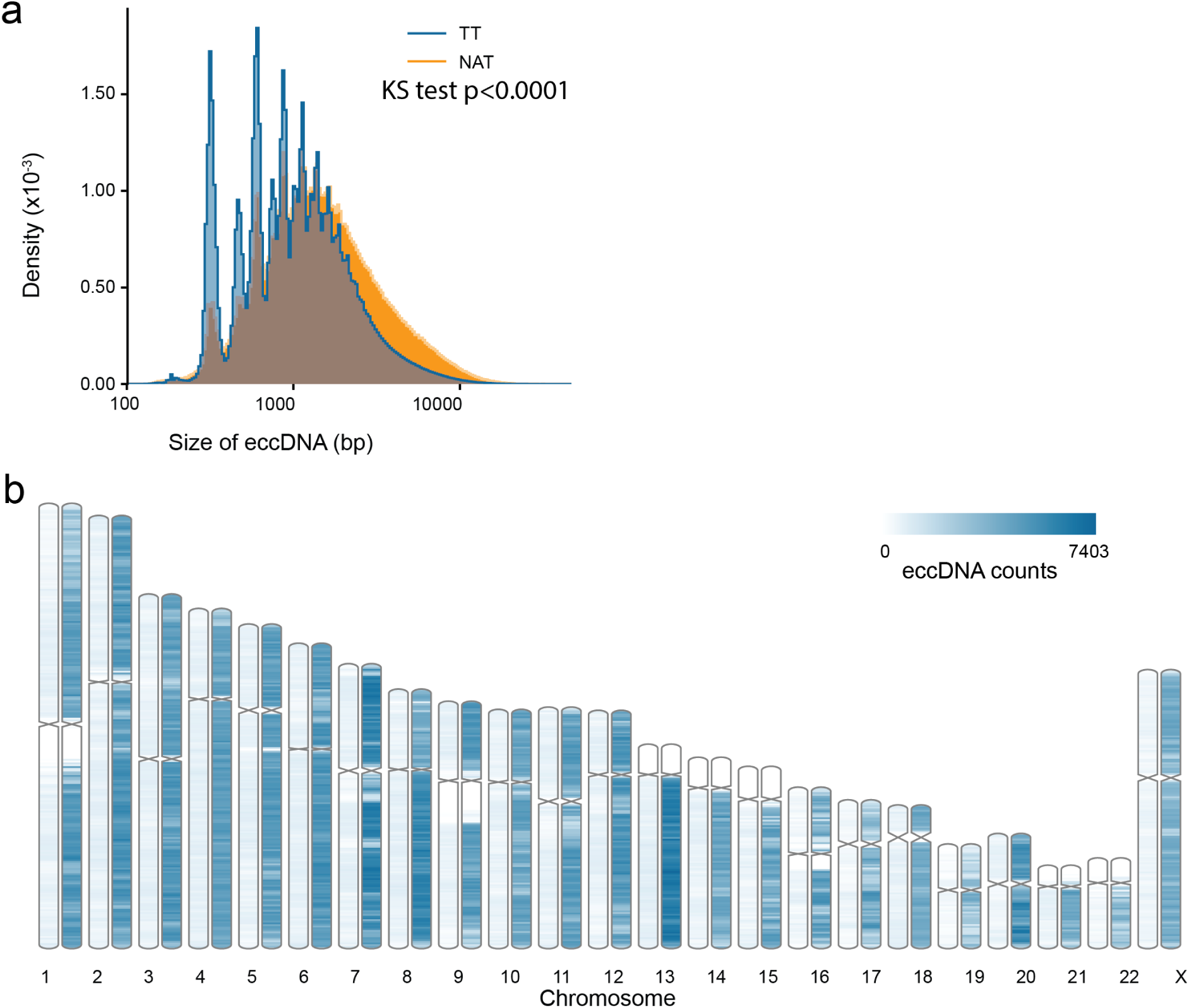
Characterization of eccDNA Profiles. **(a)** Density plot of eccDNA size distribution in TT and NAT samples. **(b)** Distribution of combined eccDNA counts across chromosomes in 25 NAT and TT samples. The bar plot displays eccDNA counts for each chromosome (1-22 and X) in NAT (left bar) and TT (right bar) samples. The x-axis represents chromosomes, while the y-axis shows eccDNA counts. The color intensity corresponds to eccDNA abundance, with darker blue indicating higher counts, as shown in the legend (0 to 7403 eccDNA counts.

**Supplementary Figure 4.**
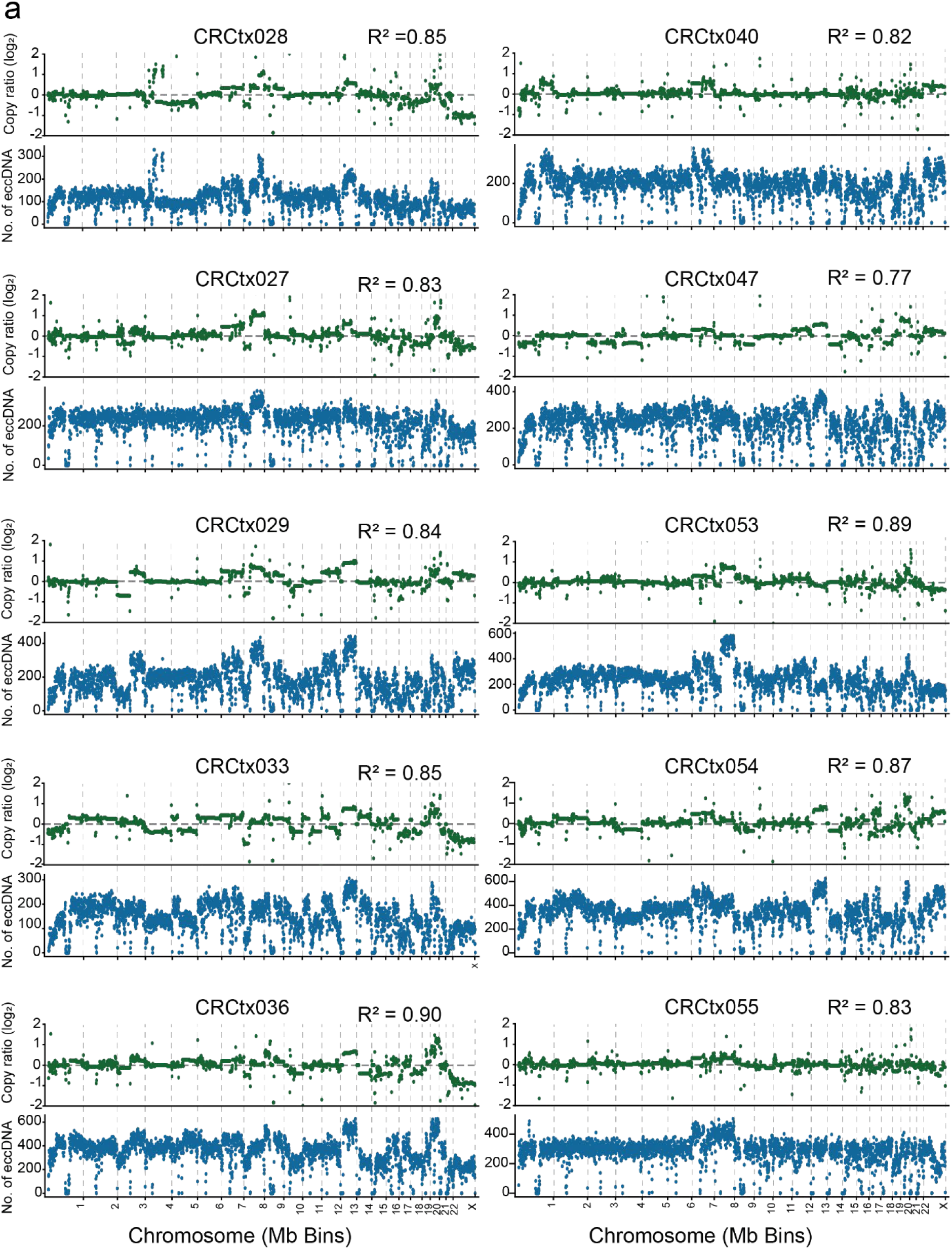
Comparison of eccDNA and copy number. **(a)** Genome wide distribution of eccDNA count per MB to the copy number ratio. The top panel (green) shows the log_2_ copy number ratio across chromosomes per MB obtained from CNVKit, while the bottom panel (blue) represents the corresponding eccDNA counts per MB for the samples in the cohort. The coefficient of determination (r^2^) determined by regression analysis is labeled in the figure.

**Supplementary Figure 5.**
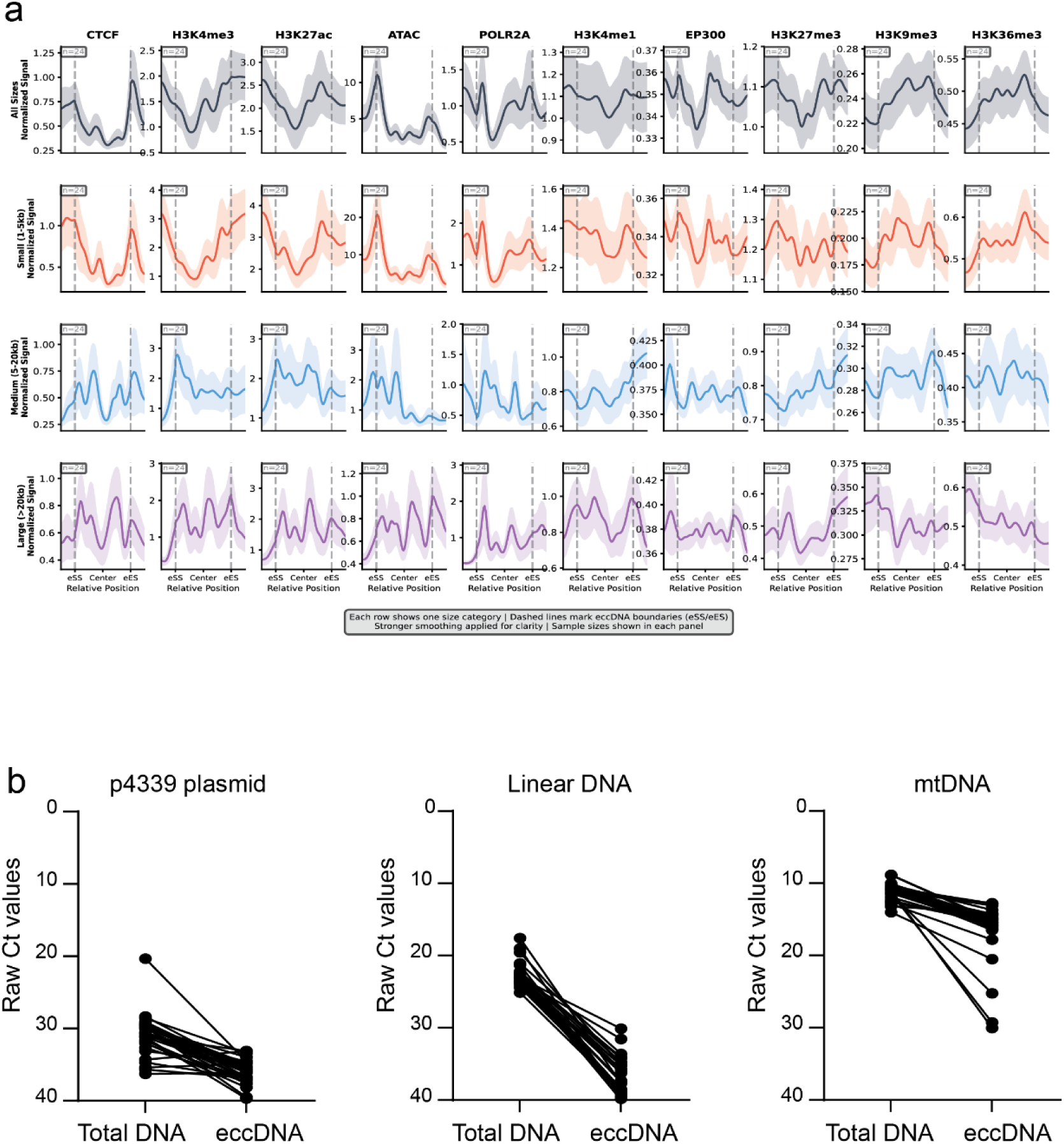
Tracks show normalized read density for: CTCF, H3K4me3, H3K27ac, ATAC, POL2A, H3K4me1, EP300, H3K27me3, and H3K36me3. **(a)** All eccDNA shown in gray tracks. **(b)** Small eccDNA (1-5kb) shown in red tracks. **(c)** Medium eccDNA (5-20kb) shown in blue tracks. **(d)** Large eccDNA (>20kb) shown in purple tracks. Each row represents one size category. Vertical dashed lines mark eccDNA boundaries (eSS, eccDNA start site; eES, eccDNA end site). X-axis represents relative position across eccDNA regions; y-axis shows normalized signal intensity (RPM). Sample sizes indicated in individual panels. Stronger smoothing applied for visual clarity. (**e)** Quality control tests for purified circular DNA were performed using a qPCR assay. Following CRISPR-Cas9 and Exonuclease V treatments, the levels of linear DNA and mtDNA in the total DNA sample were reduced. The circular p4339 plasmid, used as an internal control, remained detectable after the treatments. Ct, cycle threshold; p4339, inner control plasmid; mtDNA, mitochondrial DNA.

## Supplementary Tables

**S1: Clinical characteristics of CRC cohort**

**S2: No. of eccDNA-borne gene counts in TT, NAT and synthetic dataset. Permutation test results**

**S3: FoldChange of eccDNA-borne genes in TT with respect to NAT, binomial test results, GSEA results**

**S4: No. of eccDNA per MB in each sample and chi-square test results**

**S5: ENCODE accession numbers for ChIP and ATAC data and chromatin enrichment analysis**

**S6: Description of primers for linear DNA fragment synthesis, qPCR and PCR reactions.**

